# A horizontally acquired expansin gene increases virulence of the emerging plant pathogen *Erwinia tracheiphila*

**DOI:** 10.1101/681643

**Authors:** Jorge Rocha, Lori R. Shapiro, Roberto Kolter

**Affiliations:** Department of Microbiology, Harvard Medical School, Boston MA.; Conacyt-Centro de Investigación y Desarrollo en Agrobiotecnología Alimentaria, San Agustin Tlaxiaca, Mexico; Department of Applied Ecology, North Carolina State University, Raleigh, NC

**Keywords:** expansin, virulence, glycoside hydrolase, *Cucurbita*, *Erwinia*, squash, plant cell wall, cellulose, pectin, horizontal gene transfer, plant pathogen, xylem

## Abstract

All land plants depend on proteins called ‘expansins’ that non-enzymatically loosen structural cellulose, enabling cell wall extension during normal growth. Surprisingly, expansin genes are also present – but functionally uncharacterized – in taxonomically diverse bacteria and fungi that do not produce cellulosic cell walls. Here, we find that *Erwinia tracheiphila* (Enterobacteriaceae), the causative agent of bacterial wilt of cucurbits, has horizontally acquired an operon with a microbial expansin (*exlx*) gene and a glycoside hydrolase family 5 (*gh5*) gene. *E. tracheiphila* is an unusually virulent plant pathogen that induces systemic wilt symptoms followed by plant death, and has only recently emerged into cultivated cucurbit populations in temperate Eastern North America. Plant inoculation experiments with deletion mutants show that EXLX-GH5 is a secreted virulence factor that confers efficient xylem movement and colonization ability to *E. tracheiphila*. Bacterial colonization of xylem blocks sap flow, inducing wilt symptoms and causing plant death. Together, these results suggest that the horizontal acquisition of the *exlx-gh5* locus was likely a key step driving the recent emergence of *E. tracheiphila*. The increase in *E. tracheiphila* virulence conferred by microbial expansins, the presence of this gene in many other bacterial and fungal wilt-inducing plant pathogen species, and the amenability of microbial expansins to horizontal gene transfer suggest this gene may be an under-appreciated virulence factor in taxonomically diverse agricultural pathogens.

**Importance:** *Erwinia tracheiphila* is a bacterial plant pathogen that causes a fatal wilt infection in cucurbit crop plants. Here, we report that *E. tracheiphila* has horizontally acquired a microbial expansin gene (*exlx*) adjacent to a glycoside hydrolase family 5 (*gh5*) gene. Expansins are predominantly associated with plants due to their essential role in loosening structural cell wall cellulose during normal growth. We find that the EXLX and GH5 proteins in *E. tracheiphila* function as a single complex to facilitate xylem colonization, possibly by manipulating the size of xylem structures that normally exclude the passage of bacteria. This suggests that horizontal acquisition of the *exlx-gh5* locus was likely a key step in the recent emergence of *E. tracheiphila* as an unusually virulent plant pathogen. The presence of microbial expansin genes in diverse species of bacterial and fungal wilt-inducing pathogens suggests it may be an under-appreciated virulence factor for other microbes.

## Introduction

The surfaces of all land plants are colonized by complex microbial communities. For a microbe, the ability to colonize a plant increases access to the nutritional resources produced by that plant (1-3). This has driven the evolution of diverse molecular mechanisms for plant colonization that can be found in commensal, beneficial and pathogenic microbes (4-6). A group of particularly intriguing, yet largely uncharacterized genes identified in an increasing number of plant-associated bacterial and fungal species encode proteins called ‘expansins’ (7-10). Expansins are non-enzymatic, two-domain proteins of ∼250 amino acids; the N-terminal domain is related to glycoside hydrolase family 45 functional domains, and the C-terminal domain is related to grass pollen allergens (11-13). Expansin-coding genes are ubiquitous in all species of land plants and green algae, where they fulfill the essential role of non-enzymatically loosening cell wall cellulose during normal growth of any rapidly expanding tissues (11, 14-18). In bacteria and fungi – which do not have cellulosic cell walls – expansin genes are variably present in species that interact with live or dead plant or algal matter (7, 8, 10, 19). The function(s) of expansins in microbes are unknown for almost all species, but are thought to promote colonization of plants through interactions with plant cell wall cellulose (7, 8, 20).

In the few microbial species where expansin functions have been empirically investigated, their contributions to plant colonization have been diverse. The microbial expansin in the plant commensal *Bacillus subtilis* (BsEXLX1) has only a fraction of the *in vitro* ability to extend plant cell walls compared to plant expansins, but BsEXLX1 deletion mutants are either severely impaired, or unable to successfully colonize the surface of maize roots (20, 21). In some species of plant beneficial fungi, expansins (also referred to as ‘swollenins’) increase fungal mutualistic capabilities towards plant hosts (22, 23). Expansin function has also been investigated in several pathogens. In the bacterial plant pathogen *Ralstonia solanacearum*, an expansin deletion mutant has decreased virulence (24), and in *Clavibacter michiganensis*, studies have described contradictory expansin roles for virulence and ability to colonize xylem (24-28). Overall, fundamental questions surrounding how microbial expansins mediate plant colonization in divergent genetic backgrounds and variable ecological contexts, and the molecular mechanism(s) by which microbial expansins interact with plant structural carbohydrates remain enigmatic (7, 8, 10, 19, 20, 24, 26, 29, 30).

The bacterial plant pathogen *Erwinia tracheiphila* Smith (Enterobacteriaceae), the causative agent of bacterial wilt of cucurbits, contains an operon with an expansin gene (*exlx*) and a gene fragment with a glycoside hydrolase family 5 functional domain (*gh5*) (8, 31-33). The only geographic region where *E. tracheiphila* occurs is temperate Eastern North America, and it only infects species in two genera of cucurbit host plants: summer and winter squash (cultivars of *Cucurbita* spp.) and cucumber and muskmelon (*Cucumis sativus* and *Cucumis melo*) (34). The *E. tracheiphila* genome has undergone dramatic structural changes consistent with an evolutionarily recent emergence, including the horizontal acquisition of multiple genes likely important for virulence (34-38). Unlike most bacterial plant pathogens, *E. tracheiphila* cannot persist in environmental reservoirs, and instead is only transmitted by two species of highly specialized leaf beetle vectors (39-42). Bacterial cells can enter xylem when beetle frass containing *E. tracheiphila* is deposited near recent foliar feeding wounds or on floral nectaries (40, 43). Bacterica can then move systemically through xylem and block sap flow to induce systemic wilting, which is followed by plant death within 2-3 weeks after the first wilt symptoms appear (35, 44-46). *E. tracheiphila* costs farmers millions of dollars annually through direct yield losses and indirect control measures (34). Despite the economic burden caused by *E. tracheiphila*, no genetic determinants of bacterial pathogenesis or virulence have yet been empirically assessed.

Here, we characterize the role of the expansin-GH5 operon from *E. tracheiphila* for colonization of squash (*Cucurbita pepo*). First, we reconstruct the evolutionary histories of both the *exlx* and *gh5* open reading frames (ORFs), and find that the phylogenies of both *exlx* and *gh5* are consistent with horizontal acquisition by *E. tracheiphila*. Then, we create deletion mutants to determine the individual and combined roles of *exlx* and *gh5* for *E. tracheiphila* colonization of host plants. *In planta* inoculation experiments with the wild type and mutant strains show that these proteins are secreted, and suggest that they function as a single assembled EXLX-GH5 complex. The EXLX-GH5 protein complex is necessary for *E. tracheiphila* to efficiently colonize xylem, induce systemic wilt symptoms, and cause high rates of plant death. Together, these results suggest that the *exlx-gh5* locus as a non-canonical yet potent virulence factor, and horizontal acquisition of this locus was a key event driving the recent emergence of *E. tracheiphila* as a fatal plant pathogen that can efficiently colonize xylem. These findings highlight the continued risk of horizontal gene transfer driving an increase in pathogen virulence, and the continuing vulnerability of agricultural populations to invasion by pathogen variants or species with increased virulence.

## Results

### *Identification of a locus with an expansin gene in* Erwinia tracheiphila

A locus with two open reading frames (ORFs) flanked by mobile DNA elements was identified during manual curation of *ab initio* gene predictions in the *Erwinia tracheiphila* reference strains (31, 32) (Figure 1A). The first ORF, *Et-exlx* (AXF78871.1), is predicted to encode a protein product with 243 amino acids and both domains found in canonical expansin proteins (17, 47). The second ORF, *Et-gh5* (AXF77819.1), has 315 codons and is predicted to encode a putatively pseudogenized endo-1,4-beta-xylanase A precursor (EC 3.2.1.8) with a glycoside hydrolase family 5 (GH5) functional domain (www.CAZy.org) (48). Many *Et-gh*5 homologs in the NCBI *nr* database are between 415 – 450 amino acids, and RAST *ab initio* gene annotation predicts that the truncation to 315 amino acids eliminates cellulase activity and renders *Et-gh5* non-enzymatic (49). The sequences of both ORFs predict a signal peptide for secretion and a Signal Peptidase cleavage site (50), suggesting that the protein products are secreted and their function is extracellular.

**Figure 1.**
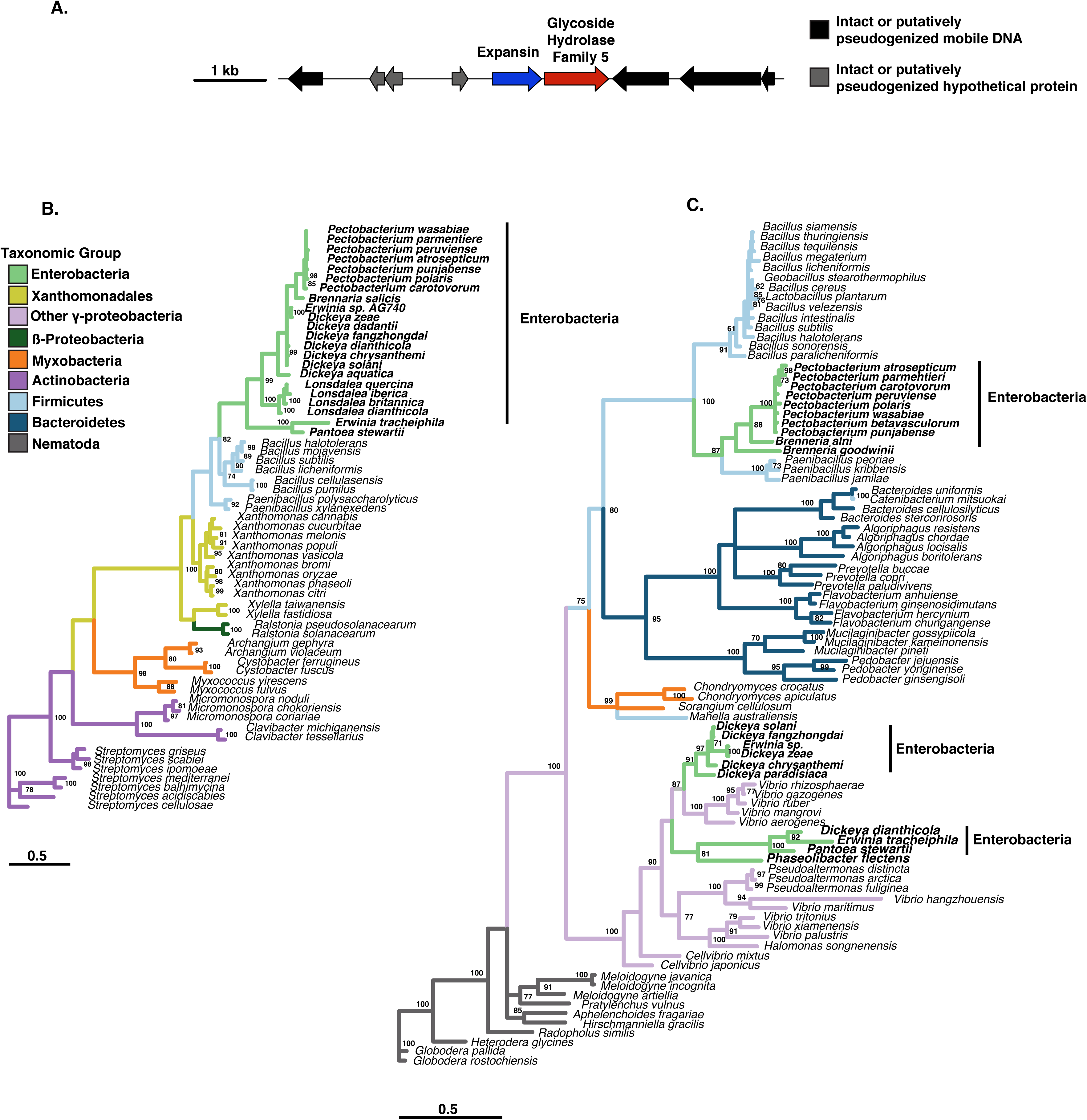
Genomic context and phylogenies of the expansin and glycoside hydrolase 5 genes in *Erwinia tracheiphila* **A)** Genomic context of the expansin (*exlx*) and glycoside hydrolase 5 (*gh5*) open reading frames (ORFs) in *Erwinia tracheiphila.* The ORFs and intergenic spaces are drawn to scale, with the black line representing position on the chromosome, and each ORF as an arrow color-coded according to *ab initio* annotated function. The scale bar is the length in nucleotides of the ORFs and intergenic spaces. **B)** Distribution of expansin (*exlx*) homologs in a taxonomically representative set of bacterial species. Branches are colored according to taxonomic assignments. The tree was reconstructed using maximum likelihood and should be considered unrooted. Numbers at nodes are bootstrap pseudoreplicate supports, and the scale bar is the number of amino acid substitutions per site. **C)** Distribution of glycoside hydrolase 5 (*gh5*) homologs in a taxonomically representative set of species. Branches are colored according to taxonomic assignments, using the same color assignments as Figure 1B. The tree was reconstructed using maximum likelihood and should be considered unrooted. Numbers at nodes are bootstrap pseudoreplicate supports, and the scale bar is the number of amino acid substitutions per site.

### *Phylogenetic distribution of the* Erwinia tracheiphila *expansin and gylocoside hydrolase family 5 open reading frames*

Because mobile DNA elements are common agents of horizontal gene transfer, the phylogenies of both the *exlx* and *gh5* genes were reconstructed and evaluated for conflict with the species phylogeny (37, 51-53). Homologs of both genes are only found in species that interact with live plants, or soil-dwelling species that likely interact with dead plant matter. The γ-proteobacterial expansin homologs are recovered as two distinct groups, separated by Firmicutes (Figure 1B). The expansin homologs in the Enterobacterial plant pathogens (*Pectobacterium* spp.*, Dickeya* spp.*, Pantoea stewartii* and *E. tracheiphila*) comprise one group, and the expansin homologs from Xanthomonadaceae comprise a second group (8). *Erwinia tracheiphila* and *Pantoea stewartii* are the only species with microbial expansin homologs from the *Erwinia* and *Pantoea* genera, respectively. This suggests that an expansin gene was horizontally acquired by an ancestral plant-associated Enterobacteriaceae species, and this original acquisition was followed by vertical and horizontal transmission between other plant-associated Enterobacteriaceae (8). The expansin phylogeny is consistent with additional horizontal gene transfer events, such as an expansin acquisition by the β-proteobacterial plant pathogen *Ralstonia solancearum* from a Xanthomonadaceae donor (8, 10).

*Gh5* homologs have a relatively sparse distribution in bacteria and some plant pathogenic nematodes, and the *gh5* phylogeny is also consistent with multiple horizontal gene transfer events (Figure 1C). *Gh5* homologs are present in the genomes of Enterobacteriaceae, Firmicutes, Myxobacteria and β-proteobacteria that also have expansin genes. Homologs are also present in species of Bacteroidetes and γ-proteobacteria species that do not have expansin genes. In Enterobacteriaceae, the *gh5* homologs separate into three distinct groups. One group is comprised of *Erwinia tracheiphila, Pantoea stewartii, Dickeya dianthicola*, and *Phaseolibacter* sp. (recently reclassified to Enterobacteriaceae (54)). The other plant-pathogenic *Dickeya* spp. comprise a second group of Enterobacterial *gh5* homologs, and plant-pathogenic *Pectobacterium* and *Brennaria* spp. are a third group.

### Distribution of expansin fusions to carbohydrate active domains in bacteria

In approximately 10% of microbial species, expansin genes are fused to domains from carbohydrate active proteins (8, 10). Out of the hundreds of families of carbohydrate active domains in the CAZy database (www.cazy.org), only GH5 and carbohydrate binding module family 2 (CBM2) domains repeatedly co-occur with bacterial expansin genes (8, 48). Bacterial species that have an expansin co-occurring with a carbohydrate active domain are more likely to be xylem-colonizing pathogens (8). The co-occurrence of GH5 domains with expansins in bacterial plant pathogens is especially apparent (Figure 2).

**Figure 2.**
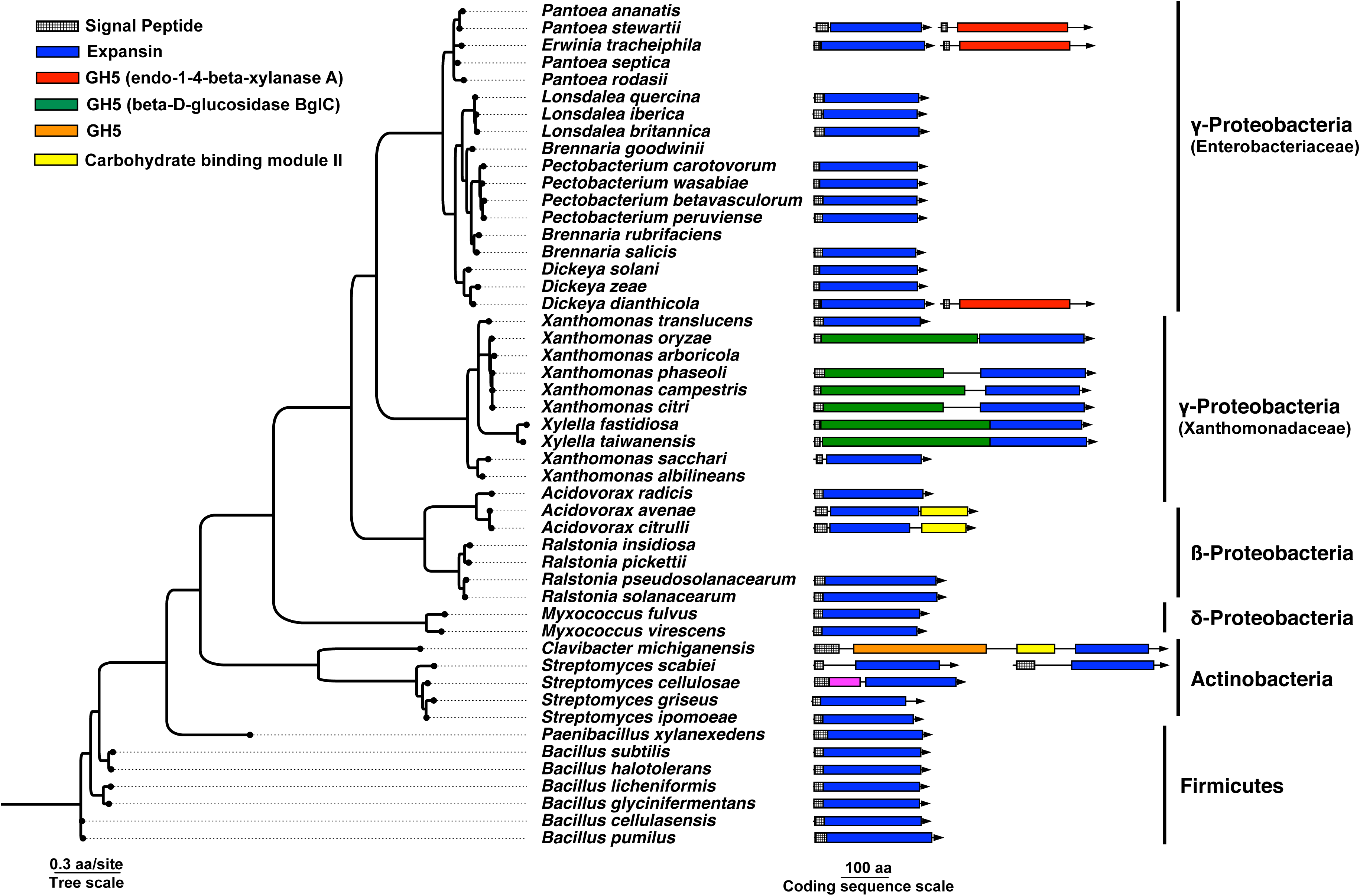
Co-occurrence of expansin genes with carbohydrate active domains. The GyrB species tree of selected bacteria with an expansin gene, and several species without expansins. The expansin and carbohydrate active domains are depicted as arrows if that species has an expansin gene, and the rectangles within the arrows indicate whether that ORF has an expansin domain, a carbohydrate active domain, or both. Homologous carbohydrate active domains are color-coded. Both expansin genes are shown for *Streptomyces scabiei*, the only microbial species to harbor two expansin homologs with signal peptides for secretion. Accession numbers of the depicted protein sequences, and accession numbers of several expansin homologs that do not have predicted signal peptides for secretion and are not depicted in the figure, can be found in Supplemental Table 3. The domains are drawn to scale. The tree was reconstructed using maximum likelihood with 100 bootstrap pseudoreplicates and should be considered unrooted.

A *gh5* homolog is present in the genomes of multiple plant-pathogenic *Pectobacterium* and *Dickeya* species that harbor an *exlx* homolog (Figure 1B, 1C). However, only three enterobacterial species (*Erwinia tracheiphila, Pantoea stewartii*, and *Dickeya dianthicola*) have the *exlx* and *gh5* ORFs in the same operon. In these three species, the *exlx* and *gh5* genes have distinct signal peptides and are separated by ∼50 nucleotides. In *P. stewartii*, the *exlx-gh5* locus is on a plasmid (pDSJ08), which may increase the probability of acting as a donor for horizontal gene transfer.

Many plant pathogenic Xanthomonadaceae have a *gh5* domain fused to an *exlx* as a single ORF (8, 55). The *gh5* domain in Enterobacteriaceae is non-homologous to the GH5 domain in Xanthomonadaceae, and the *exlx* and *gh5* domain structure in some Enterobacteriaceae is in reverse orientation compared to the *gh5-exlx* domain order in Xanthomonadaceae. A distinct *gh5* domain that is truncated to 289 amino acids is found in *Clavibacter michiganensis* (*CelA*), and this is the only known microbial expansin that is fused to both a GH5 and CBM2 domain in a single coding sequence (27). This suggests there have been at least three independent origins of an expansin adjacent or fused to a *gh5* family functional domain in bacterial plant pathogens. These multiple independent co-occurrences of bacterial expansins with evolutionarily distinct *gh5* domains may be an example of functional convergence.

### *Expansin and GH5 genes both contribute to* Erwinia tracheiphila *virulence*

The functional significance of a microbial expansin co-occuring with a carbohydrate active domain has not yet been empirically tested. To evaluate the role of EXLX and GH5 to *E. tracheiphila* virulence – and the possible synergistic effects of both proteins together – we generated a deletion mutant of the complete operon (strain Δ*exlx-gh5*), and mutants in only the expansin ORF (strain Δ*exlx*) and only the GH5 ORF (strain Δ*gh5*). Strains that complemented the three deletion mutations (Δ*exlx-gh5*(cEXLX-GH5), Δ*exlx*(cEXLX) and Δ*gh5*(cEXLX-GH5), respectively) were also constructed (Supplemental Table 1). Variation in virulence between the wild type (Wt), mutants and complemented strains were measured via squash seedling inoculation experiments. Virulence was compared by quantifying differences in the amount of time it took each strain to induce disease symptoms at three stages: 1) at initial wilt symptom development on the inoculated leaf, 2) at systemic spread of wilt symptoms to a second non-inoculated leaf, and 3) at plant death.

In plants inoculated with Δ*exlx-gh5*, wilt symptoms were delayed in the inoculated leaf and in a second systemic leaf, and significantly fewer plants inoculated with Δ*exlx-gh5* died (23%; 5 of 22) compared to plants inoculated with Wt (85%; 17 out of 20) (Figure 3A, Tables 1 and 2). Wilt symptoms in plants inoculated with Δ*exlx-gh5* were more likely to be localized to the inoculated leaf (*i.e.*, symptoms did not progress to systemic infection or plant death) compared to plants inoculated with Wt (Figure 4). The Δ*exlx-gh5*(cEXLX-GH5) complemented strain had restored ability to induce wilting symptoms at a second non-inoculated leaf, and partially restored the mortality rate (60%; 13 out of 22).

**Figure 3.**
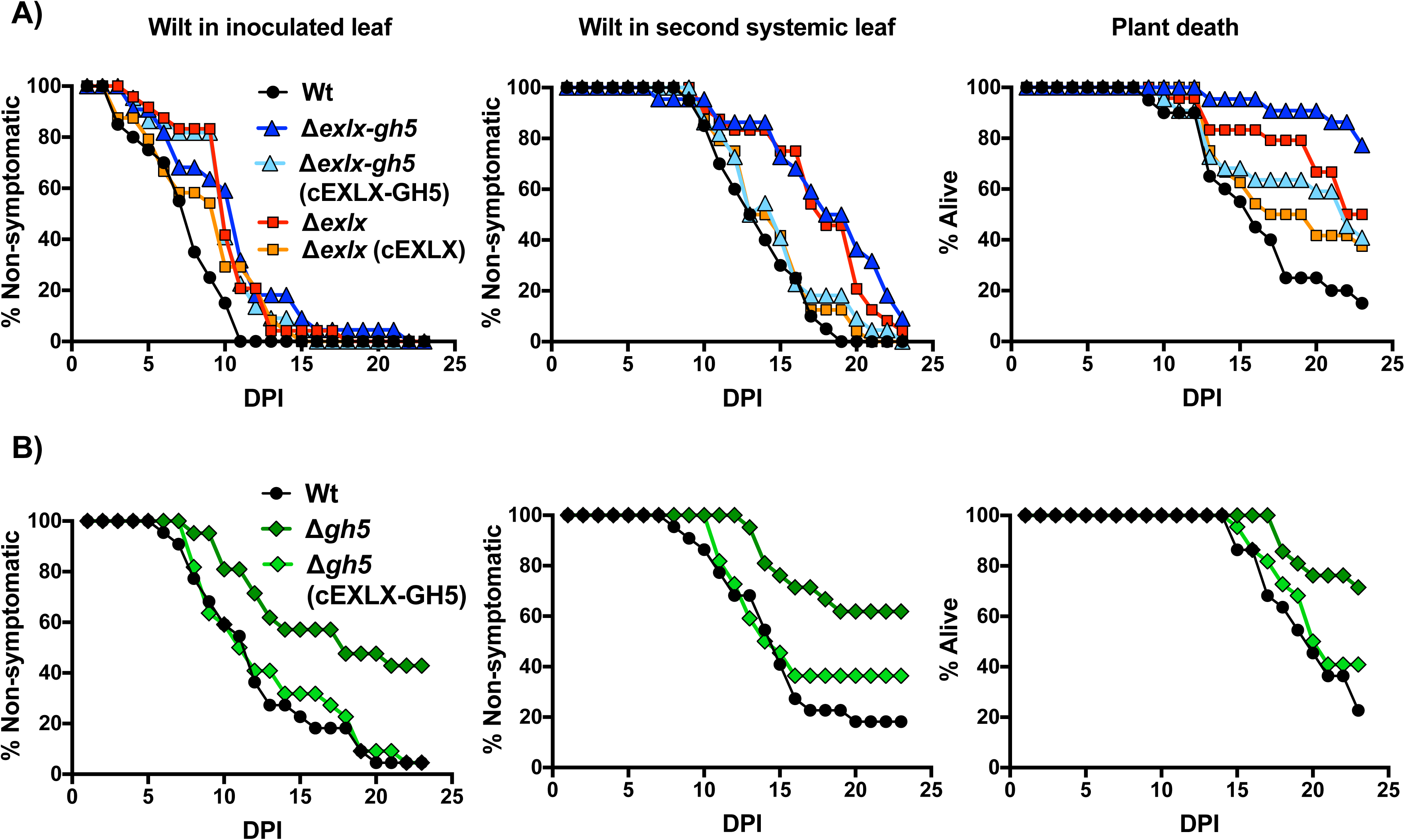
Contribution of the *Erwinia tracheiphila exlx-gh5* locus to wilt symptom development and plant death. **A)** *In planta* inoculation experiment comparing virulence of Wt, Δ*exlx-gh5*, Δ*exlx-gh5* (cEXLX-GH5), Δ*exlx* and Δ*exlx*(cEXLX) strains. **B)** A second *in planta* inoculation experiment comparing virulence of Wt, Δ*gh5*, and Δ*gh5*(cEXLX-GH5). In both **A)** and **B)**, inoculated plants were monitored for first appearance of wilt symptoms in the inoculated leaf, first appearance of systemic wilt symptoms in a second non-inoculated leaf and plant death for 23 days post inoculation (DPI). Summary and statistical analyses are in Tables 1-4. All samples sizes were between 18-22 individual plants per treatment.

**Figure 4.**
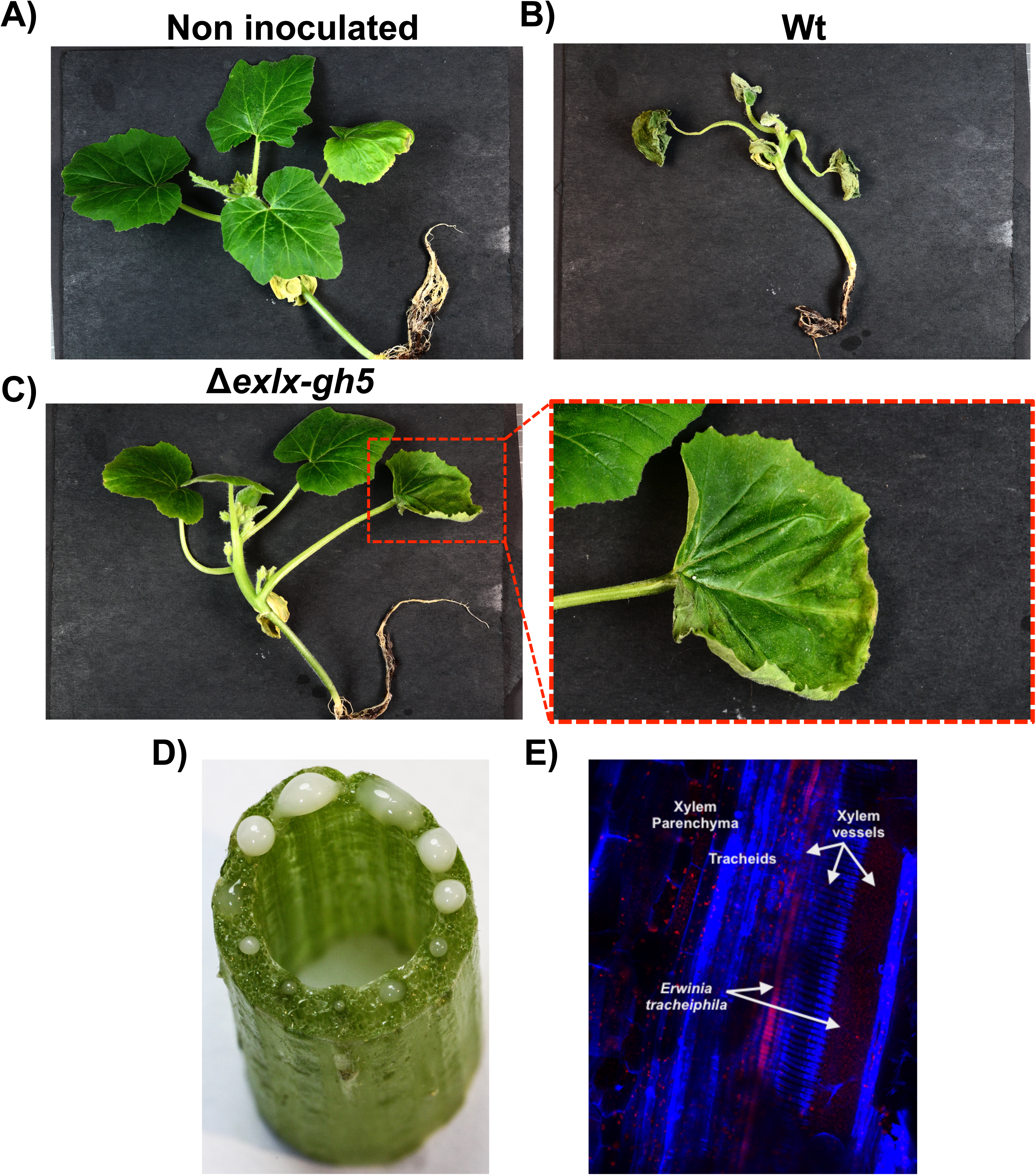
Visual comparison of wilt symptoms in squash seedlings after inoculation with either the Δ*exlx-gh5* mutant or wild type *Erwinia tracheiphila*. **A)** Non-inoculated squash seedling with no wilt symptoms **B)** Squash seedling inoculated with wild type *E. tracheiphila* that has developed systemic wilt symptoms **C)** Representative symptoms caused by inoculation with the Δ*exlx-gh5* mutant strain, where wilt often remains localized to the inoculated leaf without causing systemic wilt symptoms. **D)** Visible *E. tracheiphila* oozing from xylem in all vascular bundles of a symptomatic plant after a horizontal stem cross section cut. **E)** 20X confocal miscroscopy image of a longitudinal section of a symptomatic, *E. tracheiphila* infected *Cucurbita pepo* stem. Image is falsely colored so that plant structures are shown in blue and live *E. tracheiphila* bacterial cells are red.

**Table 1.**
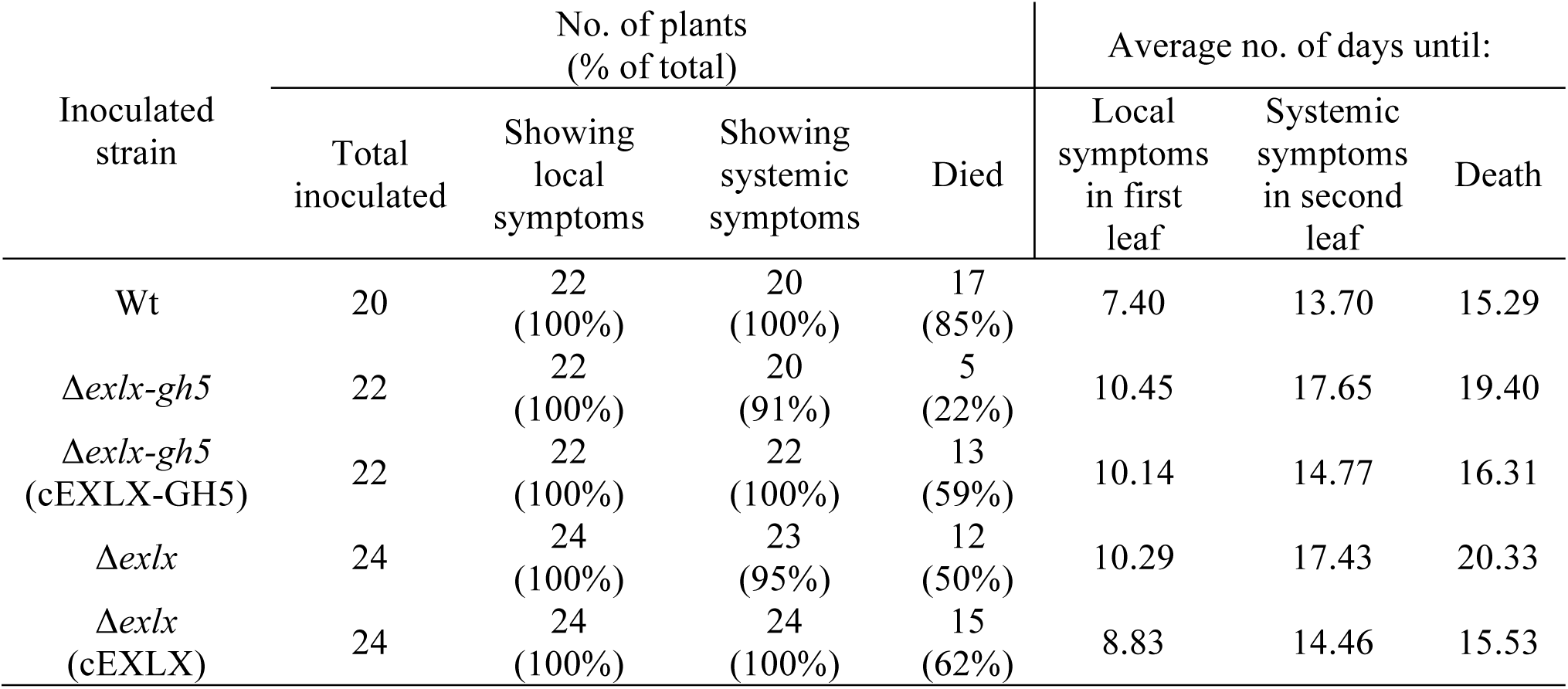
Summary of *in planta* inoculation experiment comparing virulence traits between strains Wt, Δ*exlx-gh5* mutant, Δ*exlx-gh5* (cEXLX-GH5) complemented mutant, Δ*exlx* mutant and Δ*exlx* (cEXLX) complemented mutant (corresponding to Figure 3A).

**Table 2.** Log-rank (Mantel-Cox) tests for assessing statistical differences in virulence between Wt, Δ*exlx-gh5* mutant, Δ*exlx-gh5* (cEXLX-GH5) complemented mutant, Δ*exlx* mutant and Δ*exlx* (cEXLX) complemented mutant via *in planta* inoculation experiments (corresponding to Figure 3A).

| Compared treatment groups | First leaf symptoms |  | Second systemic leaf symptoms |  | Death of plants |  |
| --- | --- | --- | --- | --- | --- | --- |
|  | Chi square | *p* value | Chi square | *p* value | Chi square | *p* value |
| All groups | 17.45 | 0.0016 | 28.51 | <0.0001 | 22.02 | 0.0002 |
| Wt vs $\Delta exlx-gh5$ | 9.787 | 0.0018 | 16.4 | <0.0001 | 20.87 | <0.0001 |
| Wt vs $\Delta exlx-gh5$ (cEXLX-GH5) | 11.04 | 0.0009 | 1.5 | 0.2206 | 3.939 | 0.0472 |
| Wt vs $\Delta exlx$ | 12.09 | 0.0005 | 16.44 | <0.0001 | 9.348 | 0.0022 |
| Wt vs $\Delta exlx$ (cEXLX) | 4.321 | 0.0376 | 0.9923 | 0.3192 | 2.497 | 0.1141 |

Individual deletions of the *Et-exlx* and *Et-gh5* ORFs also caused a decrease in virulence compared to Wt (Figure 3, Tables 1-4). Plants inoculated with either Δ*exlx* or Δ*gh5* exhibited delays in the initial appearance of wilt symptoms in the inoculated leaf, delays in the appearance of systemic wilt symptoms in a second leaf and decreased mortality compared to Wt. Genetic complementation of Δ*exlx* in strain Δ*exlx*(cEXLX) did not restore the Wt ability to cause wilt symptoms in the inoculated leaf, but did restore the ability to cause systemic wilt symptoms and plant death (Figure 3A, Tables 1 and 2).

**Table 3.** Summary of *in planta* inoculation experiment comparing virulence traits of Wt, Δ*gh5* mutant and Δ*gh5* (cEXLX-GH5) complemented mutant (corresponding to Figure 3B).

| Inoculated strain | No. of plants (% of total) |  |  |  | Average no. of days until: |  |  |
| --- | --- | --- | --- | --- | --- | --- | --- |
|  | Total inoculated | Showing local symptoms | Showing systemic symptoms | Died | Local symptoms in first leaf | Systemic symptoms in second leaf | Death |
| Wt | 22 | 22 (100%) | 18 (81%) | 17 (77%) | 11.82 | 13.61 | 18.82 |
| $\Delta gh5$ | 21 | 12 (57%) | 8 (38%) | 6 (28%) | 13.25 | 15.38 | 19.33 |
| $\Delta gh5$ (cEXLX-GH5) | 22 | 21 (95%) | 14 (66%) | 13 (59%) | 12.67 | 13.00 | 18.54 |

**Table 4.** Log-rank (Mantel-Cox) tests for assessing statistical differences in virulence between strains Wt, Δ*gh5* mutant and Δ*gh5* (cEXLX-GH5) complemented mutant (corresponding to Figure 3B).

| Compared treatment groups | First leaf symptoms |  | Second systemic leaf symptoms |  | Death of plants |  |
| --- | --- | --- | --- | --- | --- | --- |
|  | Chi | *p* value | Chi | *p* value | Chi | *p* value |

|  | square |  | square |  | square |  |
| --- | --- | --- | --- | --- | --- | --- |
| All groups | 15.53 | 0.0004 | 9.216 | 0.01 | 9.833 | 0.0073 |
| Wt vs $\Delta gh5$ | 14.11 | 0.0002 | 9.906 | 0.0016 | 10.25 | 0.0014 |
| Wt vs $\Delta gh5$ (cEXLX-GH5) | 1.059 | 0.3034 | 0.8276 | 0.363 | 1.187 | 0.276 |

Promoter regions for expression of the *exlx-gh5* operon have not been characterized, but expression of individual ORFs in an operon is often directed from an upstream shared promoter. It is therefore reasonable to assume expression of the *gh5* ORF is directed from a shared promoter region upstream of *exlx* (56). For this reason, the single Δ*gh5* mutant was complemented with the full operon (*exlx-eng*) to include the promoter region of *exlx*. Complementation of Δ*gh5* with Δ*exlx-gh5*(EXLX-GH5) restored the Wt ability to cause wilt symptoms in the inoculated leaf, systemic wilt symptoms in a second leaf, and plant death (Figure 3B Tables 3 and 4).

### The Δexlx-gh5 mutant is impaired in systemic movement

The correlation between within-plant movement of *E. tracheiphila* to systemic wilt symptom development and plant death has been hypothesized, but not yet demonstrated (34, 57). It is assumed that systemic movement of bacteria through xylem – along with bacterial replication and increase in biomass far from the initial inoculation point – is necessary to occlude xylem vessels to cause wilt symptoms and plant death (Figure 4) (34, 57). To explicitly test whether *Δexlx-gh5* has impaired within-host movement, squash seedlings were inoculated with either Wt or *Δexlx-gh5.* At 12 DPI, bacteria were quantified at two sites in the same plant: the petiole of the inoculated leaf, and the petiole of a second, non-inoculated leaf. At 12 DPI, all of the plants inoculated with the Wt strain were systemically wilting, but none of the plants inoculated with *Δexlx-gh5* had wilt symptoms beyond the inoculated leaf. At the inoculation site of all plants, the Wt and *Δexlx-gh5* both reached similar cell counts (>10^9^ CFU/g for Wt, and 10^8^-10^9^ CFU/g for *Δexlx-gh5*) (Figure 5). However, in a petiole of a second, non-inoculated leaf Δ*exlx-gh5* only reached cell counts of 10^3^ CFU/g, while the Wt reached 10^9^ CFU/g (Figure 5). The Δ*exlx-gh5* strain does not have a growth deficiency *in vitro* compared to the Wt (Supplemental Figure 1), showing that the attenuation of wilt symptom development and decrease in plant death rates (Figures 3 and 4) is due to impaired systemic movement of Δ*exlx*-*gh5* and not a difference in growth rate *per se*.

**Figure 5.**
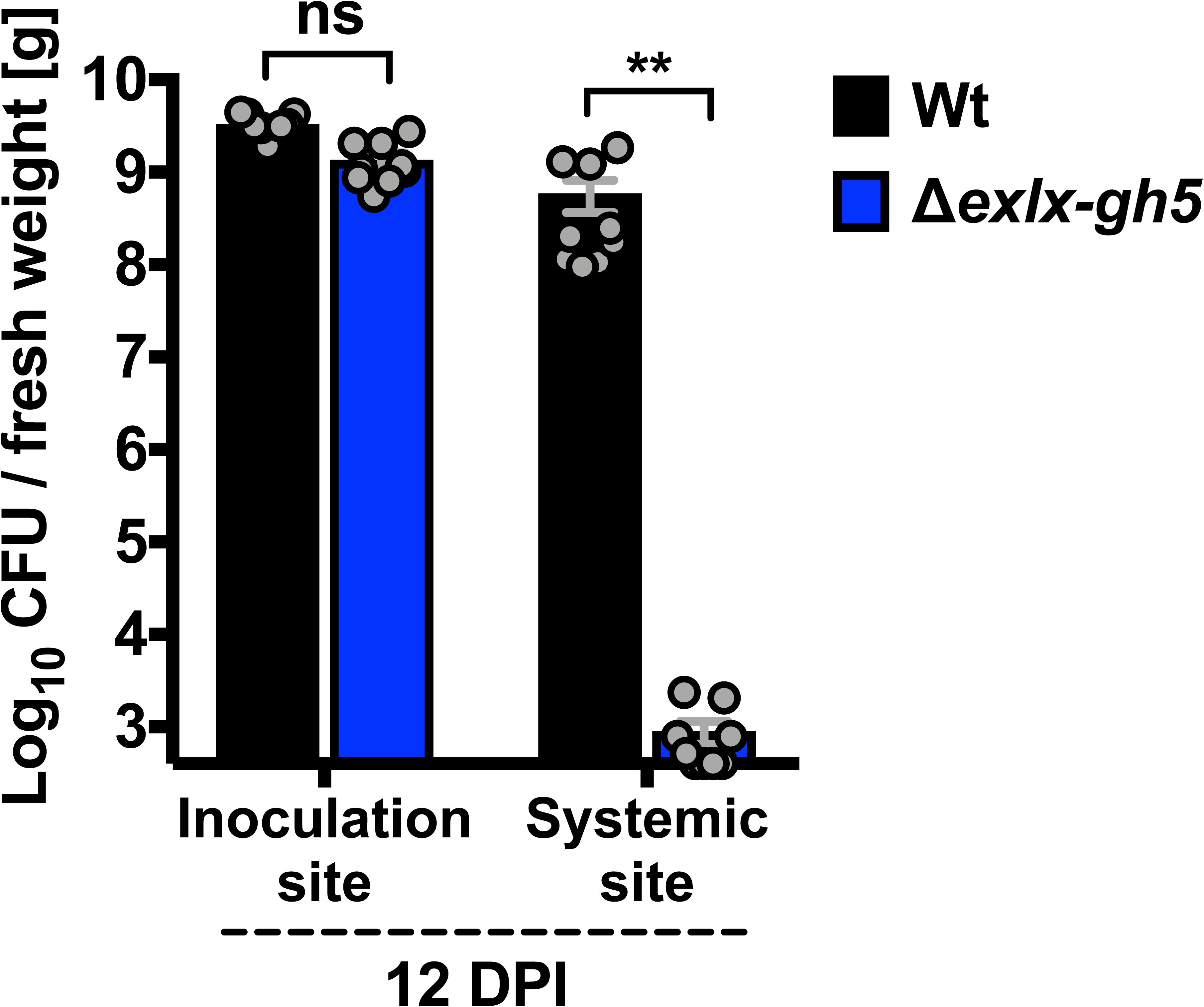
Systemic colonization capability of Wt and Δ*exlx-gh5* strains. Squash seedlings were inoculated with either Wt or Δ*exlx-gh5.* At 12 days post inoculation (DPI), bacterial concentration was determined in the inoculation site and in a petiole of a second, non-inoculated leaf. Mean ± SE are plotted, and grey circles are individual biological replicates (Sample sizes, n=9 per treatment). Y-axis is the log_10_ CFU/gram fresh weight and is scaled to the lower limit of detection for the assay (log_10_CFU/gram fresh weight = 3). Brackets indicate pairwise statistical comparisons (Dunn’s test) between two groups: ** *P* < 0.005; ns, non significant.

### *Systemic wilt symptoms are correlated with local* Erwinia tracheiphila *concentration*

A second inoculation experiment using only the *Δexlx-gh5* mutant was conducted to quantify how wilt symptom severity is correlated with the ability of *E. tracheiphila* to move through, and replicate in xylem beyond the inoculation point. Thirty plants were inoculated with *Δexlx-gh5*, and after 21 days symptoms in each plant were scored as either no symptom in any leaf (healthy), wilt symptoms restricted to the site of inoculation, systemic wilt symptoms beyond the inoculated leaf, or death. Bacteria were then quantified with CFU counts from the petiole of the inoculated leaf and a petiole of a second non-inoculated leaf.

At 21 DPI, 8 plants were healthy and had not developed any wilt symptoms, 12 only had wilt symptoms in the inoculated leaf, 10 had systemic wilting symptoms, and none had died (Figure 6). In the petiole of the inoculated leaf, the bacterial concentration reached ∼10^8^ to 10^9^ CFU/g in 26 out of the 30 experimental plants regardless of symptom severity. Cell counts from the inoculation site were lower (below <10^7^ CFU/g) in the remaining 4 plants, three of which were healthy and one that had symptoms only in the inoculated leaf (Figure 6). In the petiole of a second non-inoculated leaf, bacterial concentration was correlated with overall severity of wilt symptoms, with lower severity corresponding to lower bacterial numbers. In 6 of the 8 healthy plants that did not develop any wilt symptoms, bacterial cells were undetectable at a non-inoculated leaf, and bacterial cells in a second non-inoculated leaf of the remaining two healthy plants were just barely over the ∼ 10^3^ CFU/g threshold of detection (Figure 6). In the 12 plants that developed wilt symptoms only in the inoculated leaf, the bacterial numbers at a non-inoculated leaf were highly variable (ranging from 10^4^ to 10^8^ CFU/g). In 7 out of 10 systemically wilting plants, the bacterial concentration recovered at a second non-inoculated leaf was similar to the cell counts recovered at the local inoculation site (∼10^8^ CFU/g). These results explicitly correlate severity of wilt symptoms with the ability of *E. tracheiphila* to move systemically and increase in population far from the initial inoculation site to block xylem sap flow (Figure 4).

**Figure 6.**
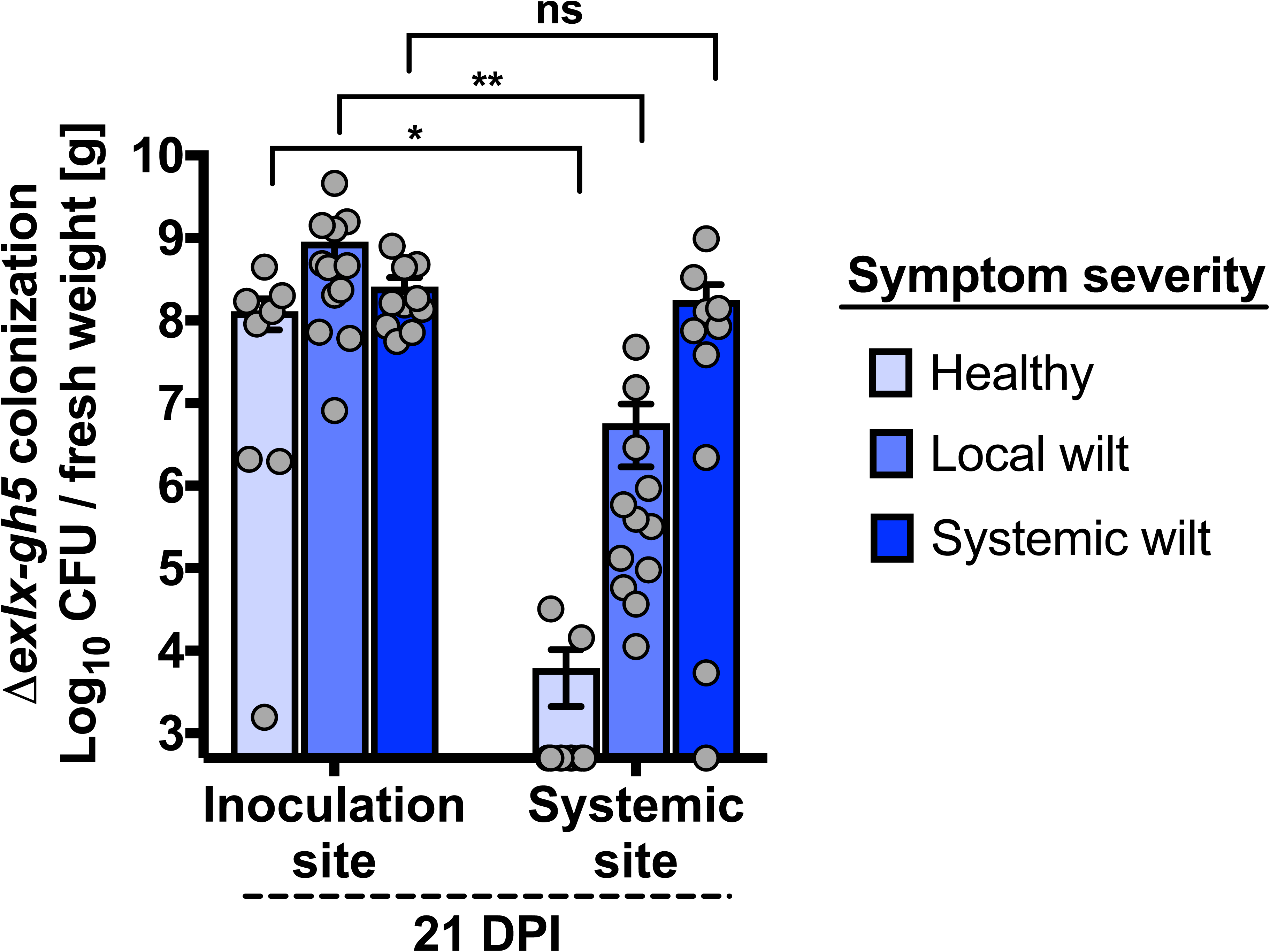
Correlation of Δ*exlx-gh5* concentration with symptom severity. Thirty squash seedlings were inoculated with Δ*exlx-gh5*, and at 21 days post inoculation (DPI) all plants were scored according to whether they remained healthy (n=8), had developed wilt symptoms only in the inoculated leaf (n= 12), or had developed systemic wilt symptoms (n=10). CFU were then counted at the inoculation site and at a second non-inoculated leaf requiring systemic movement. Bars show mean ± SE, and grey circles are individual biological replicates. Y-axis is scaled to the lower limit of detection for the assay (log_10_ CFU/gram fresh weight = 3). Brackets indicate pairwise statistical comparisons (Dunn’s test) between two groups: * *P* < 0.05; ** *P* < 0.005; ns, non significant.

### Erwinia tracheiphila *does not have cellulase or xylanase activity*

All expansin proteins (from plants, bacteria, fungi, or other microbial eukaryotes) do not have a detectable enzymatic activity (7, 13, 19, 58). However, glycoside hydrolases are enzymes that break the glycosidic bond between two or more carbohydrate subunits, and the predominant target of these enzymes is cellulose (48). It is therefore possible that the *gh5* ORF adjacent to or fused to bacterial expansin genes in some species may confer enzymatic activity. To test whether the *gh5* ORF confers carbohydrate degrading ability to *E. tracheiphila*, the Wt strain (with the intact *exlx-gh5* locus) was evaluated for enzymatic degradation of cellulose and xylan, the two main structural components of plant cell walls and the putative targets of active GH5 enzymes (59). Neither *E. tracheiphila* culture supernatant nor colonies had detectable hydrolytic activity against cellulose or xylan (Supplemental Figure 2). The absence of enzymatic activity is consistent with previous results from both plant and microbial expansins where no enzymatic activity has ever been detected (7, 13, 19).

### *Neither flagella nor Type IV Pili contribute to* Erwinia tracheiphila *systemic xylem colonization*

Type IV Pili and flagella are used by some bacterial plant pathogens during systemic movement through xylem (60-63). To assess whether these cellular components may also contribute to *E. tracheiphila* xylem colonization, deletion mutants were generated for Type IV Pili (Δ*T4P*) and flagella (Δ*fliC*) (Supplemental Table 1). In squash inoculation experiments, the virulence phenotypes of Δ*T4P* and Δ*fliC* mutants were indistinguishable from Wt. There was no difference in the development of wilt symptoms or death rate from inoculation with either Δ*T4P*, Δ*fliC*, or Wt (Figure 7, Tables 5 and 6). This indicates that neither Type IV Pili nor flagellar movement contribute to xylem colonization by *E. tracheiphila*, although it is still possible that these loci contribute in other, more subtle, ways to pathogenesis.

**Figure 7.**
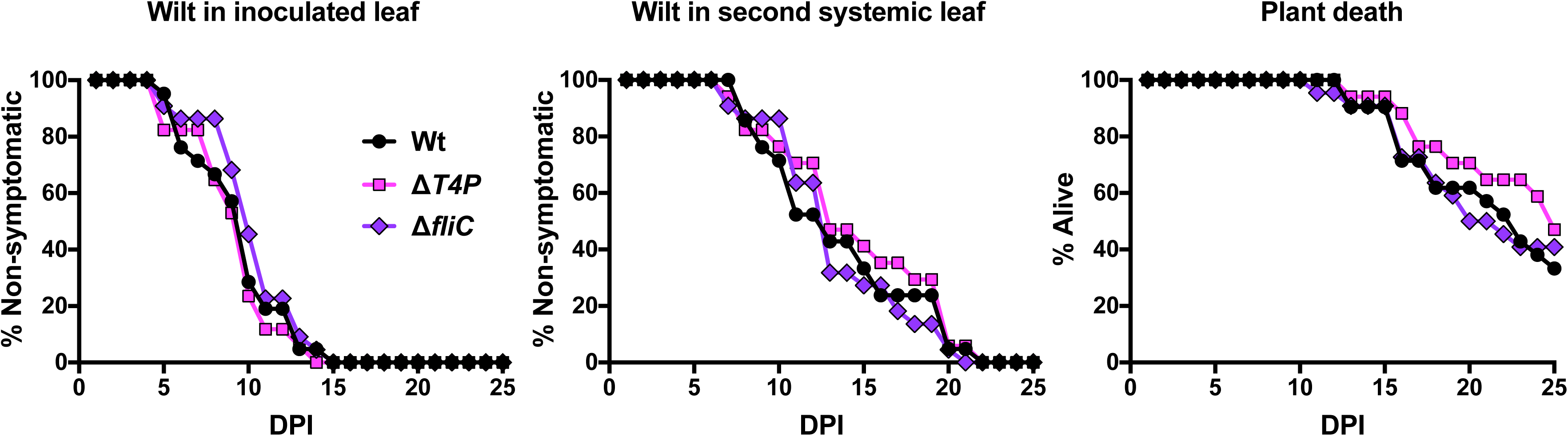
Comparison of virulence between wild type, flagellar deletion mutant and Type IV Pili deletion mutant. Squash seedlings were inoculated with either Wt, a flagellar deletion mutant (Δ*fliC*) or a Type IV Pili deletion mutant (Δ*T4P*). Inoculated plants were monitored for first appearance of wilt symptoms in the inoculated leaf, first appearance of systemic wilt symptoms in a second non-inoculated leaf and plant death for 25 days post inoculation (DPI). All samples sizes were between 17-22 individual plants per treatment. Summary and statistical analyses are in Tables 5 and 6.

**Table 5.** Summary of *in planta* inoculation experiment comparing virulence traits between strains Wt, Δ*fliC* mutant and Δ*T4P* mutant (corresponding to Figure 7).

| Inoculated strain | No. of plants (% of total) |  |  |  | Average no. of days until: |  |  |
| --- | --- | --- | --- | --- | --- | --- | --- |
|  | Total inoculated | Showing local symptoms | Showing systemic symptoms | Died | Local symptoms in first leaf | Systemic symptoms in second leaf | Death |
| Wt | 21 | 21 (100%) | 21 (100%) | 14 (66%) | 9.43 | 13.62 | 18.86 |
| $\Delta fliC$ | 22 | 22 (100%) | 22 (100%) | 13 (59%) | 10.23 | 13.45 | 17.54 |
| $\Delta T4P$ | 17 | 17 (100%) | 17 (100%) | 9 (52%) | 9.18 | 14.53 | 19.67 |

**Table 6.** Log-rank (Mantel-Cox) tests for assessing statistical differences in virulence between strains Wt, Δ*fliC* mutant and Δ*T4P* mutant (corresponding to Figure 7).

| Compared treatment groups | First leaf symptoms |  | Second systemic leaf symptoms |  | Death of plants |  |
| --- | --- | --- | --- | --- | --- | --- |
|  | Chi square | *p* value | Chi square | *p* value | Chi square | *p* value |
| All groups | 1.623 | 0.6543 | 1.324 | 0.7235 | 2.156 | 0.5406 |

### The Et-exlx-gh5 protein is secreted during infection

To test whether the protein products of the *Et-exlx-gh5* locus are secreted (as predicted by the presence of signal peptides), plants were co-inoculated with a 1:1 mix of Wt & Δ*exlx-gh5.* Successful restoration of Δ*exlx-gh5* colonization from *in trans* complementation by the Wt strain would indicate that the EXLX and GH5 proteins provided by the Wt strain are secreted and function extracellularly. An equal number of plants were inoculated with only the Wt or only the *Δexlx-gh5* mutant. In singly inoculated plants, the Wt or *Δexlx-GH5* reached the same concentration in the inoculated site at 1 DPI, but only the Wt was detected in a petiole of a non-inoculated leaf at 12 DPI (Figure 8A).

**Figure 8.**
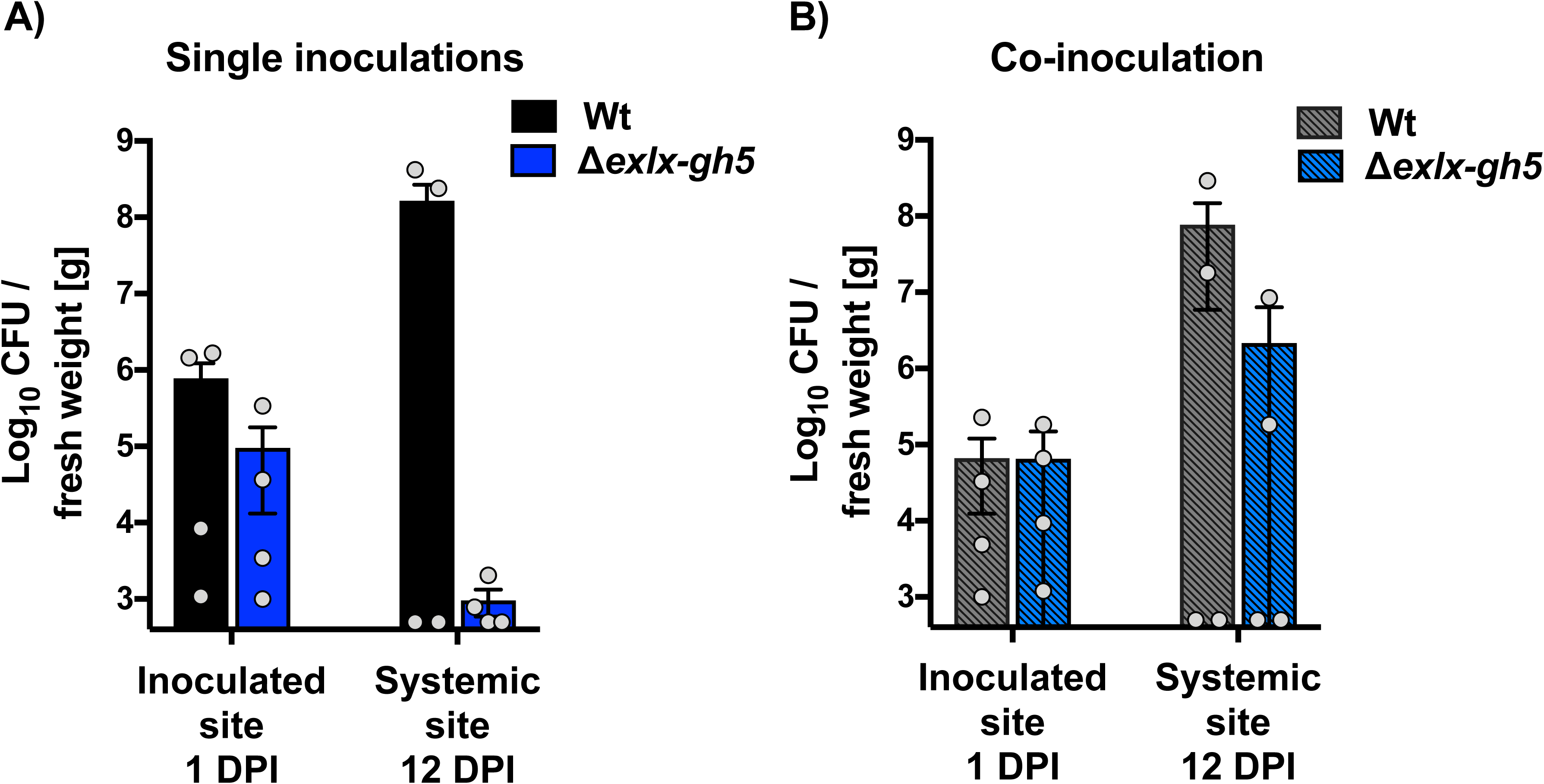
*In trans* complementation of *Δexlx-gh5* and Wt. **(A)** Plants were singly inoculated with either Wt or Δ*exlx-gh5*, and **(B)** Plants were co-inoculated with a 1:1 mix of Wt & Δ*exlx-gh5.* In both single and co-inoculation experiments, CFU were counted at 1 day post inoculation (DPI) from the local inoculation site and at 12 DPI from the petiole of a second non-inoculated leaf. Y-axis is scaled to the lower limit of detection for the assay (log_10_ CFU/gram fresh weight = 3). Bars show mean ± SE, and grey circles are individual biological replicates. Sample size, n = 4 per treatment.

In co-inoculated plants, both Wt & Δ*exlx-gh5* strains were present at similar concentrations at the inoculation site at 1 DPI (Figure 8B). After 12 days, two of the 4 plants co-inoculated with the 1:1 mix of Wt & Δ*exlx-gh5* developed systemic wilt symptoms. In these two co-inoculated plants, the cell count of Wt in the petiole of a non-inoculated leaf reached 10^7^-10^8^ CFU/g, and the cell count of Δ*exlx-gh5* reached 10^5^-10^6^ CFU/g (Figure 8B). This is a notably higher cell count than Δ*exlx-gh5* reaches at the same 12 day time point when singly inoculated (10^3^-10^4^ CFU/g) (Figure 5, Figure 8A). The ability of the Wt to partially rescue the systemic colonization defect of Δ*exlx-gh5 in trans* (when the Wt and Δ*exlx-gh5* are co-inoculated) indicates that EXLX and GH5 are secreted and function extracellularly.

To test whether the EXLX and GH5 proteins function independently or as a single assembled unit, plants were co-inoculated with a 1:1 mix of Δ*exlx* and Δ*gh5*. The Δ*exlx* deletion mutant is expected to still secrete an intact GH5 protein, and the *Δgh5* deletion mutant is expected to still secrete an intact EXLX protein. If the EXLX and GH5 proteins function independently, the two strains would complement each other *in trans*. However, co-inoculating the Δ*exlx* and *Δgh5* single deletion mutants did not rescue the attenuated virulence phenotype of the individual deletion mutants (Figure 9, Tables 7 and 8). This indicates that these proteins function as a single EXLX-GH5 protein complex that assembles before or during secretion, and therefore both proteins must be produced by the same cell.

**Figure 9.**
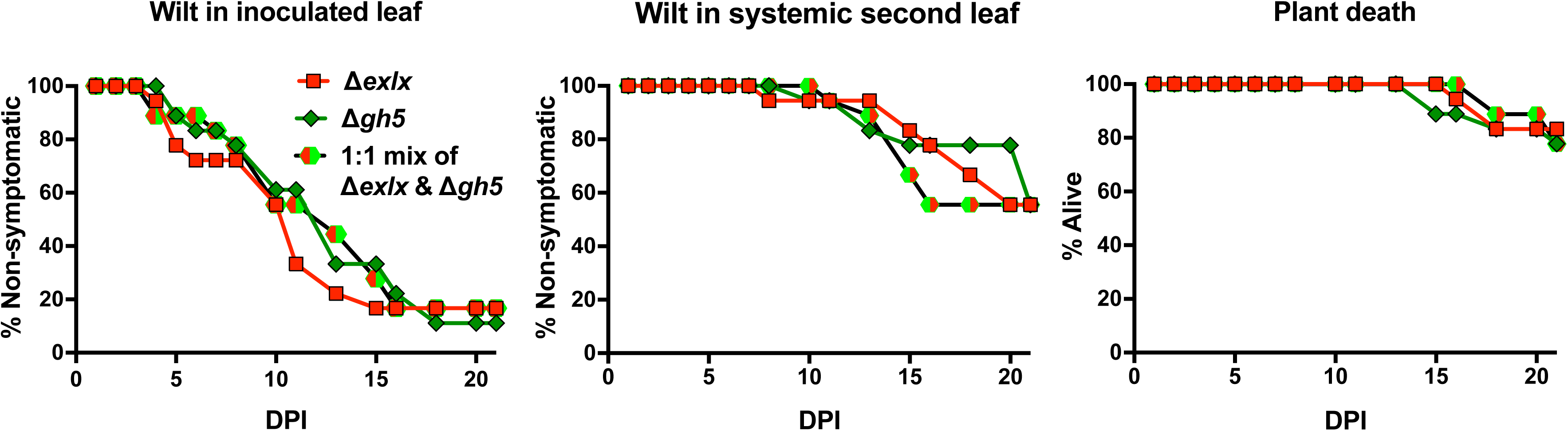
*In trans* complementation of Δ*exlx* and Δ*gh5.* Plants were co-inoculated with both the Δ*exlx* and Δ*gh5* deletion mutants. Inoculated plants were monitored for first appearance of wilt symptoms in the inoculated leaf, first appearance of systemic wilt symptoms in a second leaf and plant death for 21 days post inoculation (DPI). Sample sizes are n = 18 per treatments. Summary and statistical analyses are in Tables 7 and 8.

**Table 7.**
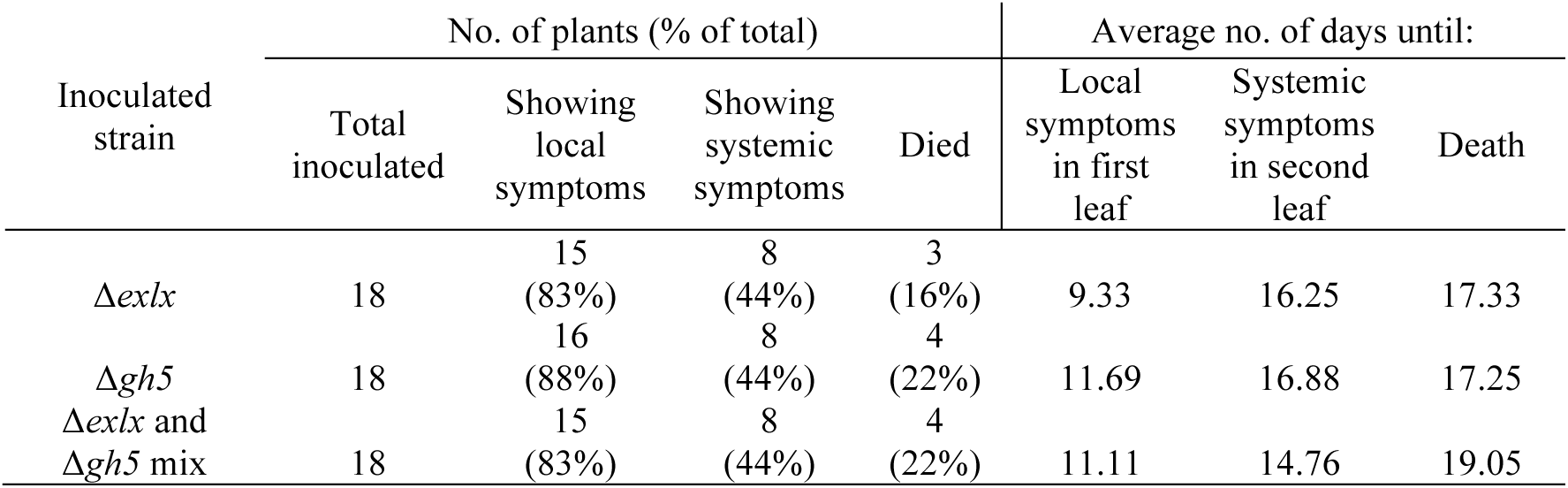
Summary of *in planta* inoculation experiment comparing virulence traits between mutant strains Δ*exlx*, Δ*gh5* and a 1:1 mix of both Δ*exlx* and Δ*gh5* mutants (corresponding to Figure 9).

| Inoculated strain | No. of plants (% of total) |  |  |  | Average no. of days until: |  |  |
| --- | --- | --- | --- | --- | --- | --- | --- |
|  | Total inoculated | Showing local symptoms | Showing systemic symptoms | Died | Local symptoms in first leaf | Systemic symptoms in second leaf | Death |
| $\Delta exlX$ | 18 | 15 (83%) | 8 (44%) | 3 (16%) | 9.33 | 16.25 | 17.33 |
| $\Delta gh5$ | 18 | 16<br>(88%) | 8<br>(44%) | 4<br>(22%) | 11.69 | 16.88 | 17.25 |
| $\Delta exlx$ and<br>$\Delta gh5$ mix | 18 | 15<br>(83%) | 8<br>(44%) | 4<br>(22%) | 11.11 | 14.76 | 19.05 |

**Table 8.** Log-rank (Mantel-Cox) statistical test results for experiment comparing virulence of mutant strains Δ*exlx*, Δ*gh5* and a 1:1 mix of both Δ*exlx* and *Δgh5* mutants (corresponding to Figure 9).

| Compared treatment groups | First leaf symptoms |  | Second systemic leaf symptoms |  | Death of plants |  |
| --- | --- | --- | --- | --- | --- | --- |
|  | Chi square | *p* value | Chi square | *p* value | Chi square | *p* value |
| All groups | 0.5335 | 0.7659 | 0.05808 | 0.9714 | 0.2023 | 0.9038 |

## Discussion

Here, we find that the emerging plant pathogen *Erwinia tracheiphila* has horizontally acquired an *exlx-gh5* locus that functions as a virulence factor by conferring the ability to systemically colonize xylem, block sap flow and cause high rates of plant death. The ability of a pathogen to move systemically through host vasculature – either plant xylem or animal cardiovascular systems – is a high-virulence phenotype, and is associated with development of more severe symptoms than localized infections (64, 65). The ability of a pathogen to reach a high titre and be distributed throughout the host’s vasculature is also necessary for vector transmission by providing more opportunities for acquisition (64, 66). In *E. tracheiphila*, the development of systemic wilt symptoms induces a chemical volatile phenotype that attracts significantly more foraging vectors to wilting leaves (45, 67), and a physical phenotype that facilitates insect vector feeding – and increased pathogen acquisition opportunities – from symptomatic foliage (40, 45). This increase in virulence conferred by the *Et-exlx-gh5* locus induces more severe symptoms in infected plants that both attract obligate insect vectors to infected plants, and facilitates preferential feeding on wilting tissue once they arrive. Together, this suggests that the horizontal acquisition of the *exlx-gh5* locus was a key step in the recent emergence of *E. tracheiphila* as a virulent wilt-inducing pathogen that is obligately insect vector transmitted (34, 37).

In ∼10% of bacterial species that have expansin genes, the expansin is fused to domains from carbohydrate active proteins. The formation of new genes via fusions of multiple modular domains is a key source of evolutionary innovation for organisms across the tree of life (68, 69). Expansin co-occurrence with *gh5* domains is over-represented in pathogenic bacterial species that can move through xylem (8), suggesting there may be emergent properties of the EXLX-GH5 protein complex that are uniquely adaptive for xylem-colonizing plant pathogenic bacteria. Healthy plants have effective physical barriers to allow the flow of xylem sap while excluding bacteria; pit membranes between adjacent tracheids and perforation plates between xylem vessels are openings on a nanometric scale, while most bacteria are ∼1µm (70). One hypothesis is that bacterial expansins may non-enzymatically ‘loosen’ the cellulose and pectin matrix at the perforation plates or at the pit membranes in order to increase their size enough to allow the passage of bacterial cells (71-74). The ability of *E. tracheiphila* Wt strain to complement Δ*exlx-gh5 in trans* is consistent with the hypothesis that the EXLX-GH5 protein complex functions extracellularly by interacting with xylem structrual carbohydrates that would normally prevent bacterial passage. This also suggests that, while the *Et-gh5* enzymatic activity has been lost due to truncation, the remaining fragment may have been neofunctionalized and is providing an essential (though mechanistically undefined) role in virulence. One possibility is that the *gh5* functional domain may physically (but non-enzymatically) interact with plant structural carbohydrates at perforation plates or pit membranes in a way that aids expansin function for loosening of cellulose microfibrils, or *vice versa*.

The distinct phylogenies of the *Et-exlx* and *Et-gh5* ORFs, and their genomic architecture as distinct genes in the same operon, may offer mechanistic insight into how bacterial expansins fuse to carbohydrate active domains. In many Firmicutes, *Pectobacterium* spp. and most *Dickeya* spp. plant pathogens, the *exlx* and *gh5* homologs are present in the same genome, but are not located directly adjacent to each other in the same operon. Only in *E. tracheiphila, P. stewartii*, and *D. dadantii* is the *exlx* homolog directly adjacent – but not fused to – the *gh5* homolog. This suggests that during a horizontal gene transfer event between an Enterobacteriaceae donor and recipient, an expansin integrated by random chance adjacent to a GH5, and the two ORFs in this operon are now being horizontally transferred together. The assembled protein complex produced by the *exlx-gh5* locus may provide a more efficient mode of action for movement through xylem, promoting the fitness of the host bacteria and providing opportunities for further horizontal transfer of this construct as a single virulence island. From a shared promoter and only ∼50 nucleotide separation, a fusion of *exlx* and *gh5* into a single ORF is possible from a small deletion mutation. We also note that all three of the bacterial plant pathogens with this construct are agricultural pathogens emerging into intensively cultivated, homogeneous crop plant populations. *Erwinia tracheiphila* has recently emerged into cucurbit agricultural populations (34, 37) and *Pantoea stewartii* infects sweet corn (75). Both of these pathogen species only occur in temperate Eastern North America – one of the world’s most intensively cultivated regions – despite global distrubtion of susceptible host plants (76). *D. dianthicola* causes a virulent wilt disease and is emerging into cultivated potato crops, and is also geographically restricted to Eastern North America and Europe (77-79).

There is constant risk that agro-ecosystems will be invaded by virulent microorganisms, and the increasing homogeneity in crop plant populations may select for novel pathogens with non-canonical virulence mechanisms. The recent realization that microbial expansin genes are present in phylogenetically diverse xylem-colonizing bacterial and fungal species – including almost all of the most economically damaging bacterial and fungal wilt pathogens – and the function of expansins to increase *E. tracheiphila* virulence suggest these genes may be an under-appreciated virulence factor (7, 8, 80, 81). The emergence of virulent plant pathogens that systemically colonize xylem is especially alarming because plants do not have inherent genetic resistance against xylem-dwelling vascular pathogens (82). That expansin genes can confer an increase in pathogen virulence, are present in many damaging wilt-inducing agricultural plant pathogens, and are amenable to horizontal gene transfer should raise concerns about whether this gene is a more important virulence factor of agricultural plant pathogens than currently recognized.

## Methods

### Study System

*E. tracheiphila* is one of the few plant pathogenic bacterial species that moves systemically through xylem and causes host death, compared to most species that cause localized foliar lesions (83). *E. tracheiphila* is obligately vector-transmitted by two species of highly specialized leaf beetles, the striped cucumber beetle *Acalymma vittatum* and the spotted cucumber beetle *Diabrotica undecimpunctata howardii* (Coleoptera: Chrysomelidae: Luperini: Diabroticina). These herbivores have co-evolved with wild *Cucurbita* spp. in the New World, and are among the only herbivores that can detoxify ‘cucurbitacins’ (oxygenated tetracyclic triterpene), which are a class of secondary metabolites produced by many Cucurbitaceae (35, 84, 85). The striped cucumber beetle (*Acalymma vittatum*), which is obligately dependent on *Cucurbita* in all life stages, is the predominant vector species driving *E. tracheiphila* epidemics (40, 67, 86). Striped cucumber beetles acquire *E. tracheiphila* by feeding on wilting, infective foliage, which is physically easier for them to consume than non-wilting foliage (36, 45). *E. tracheiphila* colonizes the beetle hindgut (40, 87-89), and beetles can transmit *E. tracheiphila* when frass (poop) from infective beetles is deposited near recent feeding wounds on foliage, or on floral nectaries (36, 40, 43, 86). The only known overwinter reservoir for *E. tracheiphila* is infective cucumber beetles, which diapause as adults and infect new seedlings in early spring when they emerge (39, 40, 45, 86, 89, 90).

*Erwinia tracheiphila* is an example of a plant pathogen that has recently emerged into a new ecological niche created by construction of homogeneous agro-ecosystems (34, 35, 37). Analysis of the *E. tracheiphila* genome shows this species has undergone among the most dramatic structural genomic changes of any bacterial pathogen, of any host species. These changes include genome decay through pseudogenization, invasion and proliferation of mobile genetic elements, and horizontal gene acquisitions. Together, these are the canonical genomic signatures of a recent specialization on a new host species or population (34, 37, 38). *Erwinia tracheiphila* only infects few species in two genera of the cosmopolitan plant family Cucurbitaceae. One of the genera that suffers economic losses, *Cucurbita* spp. (squash, pumpkin, zucchini and some gourds), are native to the New World tropics and subtropics (91, 92). Two *Cucumis* spp. (cucumber *Cucumis sativus;* and muskmelon *Cucumis melo*) native to the Old World tropics and subtropics are the most susceptible hosts to *E. tracheiphila* infection, and the introduction of highly susceptible *Cucumis* crop plants into temperate Eastern North America likely drove the recent emergence of this pathogen (93-95). While these susceptible cucurbit cultivars are among the highest acreage crop plants globally (http://www.fao.org/faostat/), *E. tracheiphila* only occurs in temperate Eastern North America (34, 96).

### Bacterial strains, culture media and plant cultivation

All bacterial strains used in this study are listed in Supplemental Table 1. Throughout this work, we used a rifampicin resistant variant of *Erwinia tracheiphila* BHKYR (Wt) (34). *Escherichia coli* TOP10 and PIR1 strains for used for routine cloning, and the *E. coli* strain S17-1λ was used as the donor for conjugation. *E. tracheiphila* was grown in KB liquid media or agar at room temperature (RT), and *E. coli* strains in LB media or agar at 37°C, unless otherwise specified. Antibiotics were added to liquid or agar media at the following concentrations: rifampicin, 50 µg/ml; ampicillin or carbenicillin, 100 µg/ml; chloramphenicol 5 µg/ml; kanamycin 50 µg/ml. All *in planta* experiments were conducted with organic ‘Dixie’ variety crookneck squash bought from Johnny’s Seeds (https://www.johnnyseeds.com/). Plants were grown in potting mix in standard six cell seedling trays in a greenhouse environment set to 25°C, 70% humidity, and a 12 hr day: 12hr night light cycle.

### *Visualization of fluorescent* Erwinia tracheiphila *in wilting squash seedlings*

*E. tracheiphila* BuffGH was transformed with a plasmid carrying the mCherry gene for visualization of fluorescent cells in symptomatic squash seedlings. Competent *E. tracheiphila* were prepared as described previously (34, 97). Briefly, cells were prepared by growing *E. tracheiphila* to an OD_600_ of 0.02. Cells were then washed with decreasing volumes, once with chilled sterile Milli-Q water and twice with 10% glycerol, and resuspended in 1/100 volume of chilled 10% glycerol. Plasmid pMP7605 was used for electroporation in a 0.2-cm cuvette, at 2.5 kV for 5.2 to 5.8 ms. Cells were incubated at room temperature without shaking for 1 h in 3 ml KB liquid and then plated in KB agar with ampicillin. Colonies of fluorescent *E. tracheiphila Et* (pMP605) were obtained after 5 days at room temperature. Ten µl of a *Et* (pMP7605) stationary culture were used for inoculating two week-old squash seedlings (at the two leaf stage), and confocal microscopic observations were performed once symptoms appear using fresh longitudinal cuts of the inoculated petiole.

### Phylogenetic reconstruction of the expansin and endoglucanase genes and comparison of domain architecture

The amino acid sequences of the expansin (WP_046372116.1) and *gh5* (WP_016193008.1) ORFs in the *Erwinia tracheiphila* reference strain (31) were used as queries to identify expansin and *gh5* homologs using the BLASTP web interface (98). A taxonomically representative sample of the top BLASTP hits for each gene were aligned using MAFFT v. 7.305b and default parameters (99). The expansin alignment was trimmed visually such that the two canonical expansin domains were conserved in the alignment, and the *gh5* alignment was trimmed with trimAI using the –automated 1 option (100). For both alignments ProtTest v. 3.4.2 was used to identify the best-fitting substitution model by BIC score, which was WAG+G for the expansin gene alignment and LG+I+G for the GH5 alignment (101). The GyrB species tree was constructed by using the *E. tracheiphila* GyrB sequence (KKF36621.1) as a query on the BLASTP web interface (98). The GyrB amino acid sequences from species known to have an expansin gene or an expansin fusion to a domain from a carbohydrate active protein were downloaded and added to a multi-fasta. The GyrB sequences were aligned with MAFFT v. 7.305b and default parameters (99).

Phylogenetic trees were reconstructed using maximum likelihood with RAxML (102) and the appropriate evolutionary model on the CIPRES server (103). The expansin tree was reconstructed with 1000 bootstrap pseudoreplicates, and the GH5 and GyrB trees were reconstructed with 100 bootstrap pseudoreplicates. The bootstrapped pseudosamples were summarized with SumTrees v. 4.4.0 (104). The resulting phylogeny was visualized in the R statistical environment using the ggtree library (105, 106). Amino acid sequences were analyzed with NCBI CBD tool to identify domain architecture (47), and signal peptides were predicted with SignalP (50). The genomic context of the *Et-exlx-gh5* locus was visualized with genoPlotR (107). Alignment files and phylogenetic scripts are available at https://github.com/lshapiro31/gh5.expansin.phylogenetics.

### Construction of deletion mutants

Mutants with a deletion in the *exlx-gh5* operon, *exlx* gene, *gh5* gene, the Type IV pili operon and the *fli*C gene were generated from an *E. tracheiphila* isolate BHKYR parental strain by double homologous recombination, using the suicide plasmid pDS132 (108). This plasmid was improved by inserting in the *Xba*I site, a constitutive *mCherry* gene amplified from plasmid pMP7605 (109) using primers JR72 and JR73 (Supplemental Table 2). The resulting plasmid (pJR74, Supplemental Table 1) allows rapid screening of conjugants colonies and colonies that have lost the plasmid. For the target genomic region to create each mutant, regions upstream of the target locus were amplified with primers pair F5 and R5, and downstream regions were amplified with primer pair F3 and R3 (See Supplemental Table 2 for specific primer names and sequences). An ampicillin resistance *bla* gene, coding for Beta-lactamase was amplified from pDK46 (97) using primers LS23 and LS24. Constructions consisting on each upstream and downstream region flanking the *bla* gene were used for *exlx-gh5*, *gh5*, *fliC* and Type 4 Pili mutants, while a construction with no flanked antibiotic cassette was prepared for the *exlx* deletion. All constructions were assembled using the Gibson Assembly Master Mix (New England Biolabs, Ipswich, MA), and then each was reamplified with nested primers containing *Sac*I restriction site (primers *Sac*I-F and *Sac*I-R, Supplemental Table 2). Constructions for *exlx-gh5*, *exlx*, *gh5*, *fliC* and Type 4 Pili deletion were inserted into the *Sac*I site of plasmid pJR74, obtaining plasmids pJR150, pJR323, pJR324, pJR74a and pJR149, respectively (Supplemental Table 1). These plasmids were transformed into *Ec*-PIR1 for preservation, and then into *Ec*-S17 for conjugation using *E. tracheiphila* as recipient. MCherry fluorescent *E. tracheiphila* conjugants were obtained in KB agar with rifampicin and chloramphenicol, then a few colonies were picked and grown in 3 ml liquid KB with chloramphenicol to stationary phase, and 100 µl were spread in KB agar with 5% sucrose and carbenicillin (or no antibiotic in the case of *exlx* deletion). Non-fluorescent, chloramphenicol-sensitive colonies were picked, PCR checked for the correct deletion and cryogenically stored in 15% glycerol at −80C.

### Genetic complementation

A new integration plasmid, specific for a neutral region in the chromosome of *Et*-BHKYR, was constructed from plasmid pJR74. To create this plasmid, two ≈0.8 Kb adjacent DNA fragments were PCR amplified form *Et*-BHKYR genomic DNA using primer pairs JR143-JR144, and JR145-JR146 (Supplemental Table 2). These fragments were ligated using the Gibson Assembly Master Mix (New England Biolabs, Ipswich MA), and reamplified using primers JR143 and JR146. The ≈1.6 Kb product was inserted in the SacI site of pJR74, generating plasmid pJR315 (Supplemental Table 1). Single cutting *Xho*I and *Bgl*II sites were engineered in the middle of the amplified neutral regions, which can be used for the insertion of complementation genes. For the complementation of the *exlx-gh5* locus or the individual *exlx* gene, the genomic region together with its natural promoter were amplified from *E. tracheiphila* genomic DNA using primers JR152 and JR154, or JR152 and JR153 (Supplemental Table 2) respectively, and DNA products were inserted into the *Xho*I site of pJR315. Each resulting plasmid was transformed into *Ec*-S17-1λ cells, which were then used as donors for conjugation with mutant strains Δ*exlx-gh5*, Δ*exlx* or Δ*gh5* as recipients. Conjugant colonies were used for negative selection with sucrose, as described above, and colonies carrying the *exlx-gh5* operon or the *exlx* gene integrated in the expected site were confirmed by PCR.

### In planta inoculation experiments

*In planta* virulence assays were performed by inoculating squash seedlings with *E. tracheiphila* Wt and derived strains, and monitoring wilt symptom development for approximately three weeks (between 21-25 days per experiment). To create inoculum, one bacterial colony of each strain was picked and added to 3 ml of liquid KB media with the appropriate antibiotic, and grown with shaking for 24 h. Then, 2-3 week-old squash seedlings (2-3 true leaves) were inoculated by manually inducing a wound where xylem was exposed in the petiole at the base of the first true leaf and adding 10 µl of culture containing ≈ 1×10^7^ bacterial cells directly into the wound. Plants were kept at 25°C, 70% humidity, and a 12 hr day: 12hr night light cycle 25C, and monitored daily for appearance of first symptoms in the inoculated leaf, appearance of wilt symptoms in a second non-inoculated leaf, and plant death.

### In planta *Colony Forming Unit (CFU) counts*

Bacterial colony forming units (CFU) counts were determined from plants inoculated with *Et-*BHKY and derived strains. Bacterial cells can be obtained directly from petioles of infected plants (Figure 4D). Two cm samples of the petiole from the inoculated leaf, or from a second non-inoculated leaf were cut from the plants and washed briefly with 70% ethanol (EtOH). Excess EtOH was removed with a paper towel and petioles were surface-sterilized over a gas flame for 1 second and placed in a sterile plastic petri dish. From each petiole sample, 10-15 disks small disks (<1 mm) were manually cut with a sterile blade and collected in a 2 ml microtube. The weight of each 2ml tube with all leaf disks was recorded to be used for normalizing CFU per gram of plant tissue in each sample, and 500 µl of chilled PBS was added to each tube. After an incubation of 40 min on ice (vortexing every 10 min) 200 µl of PBS from each tube was pipetted into a new microtube and used for serial dilutions and plating onto KB agar with rifampicin. Bacterial CFU per gram of fresh tissue was calculated.

For obtaining CFUs of individual strains in plants co-inoculated with *E. tracheiphila* Wt and Δ*exlx-gh5* mutant, serial dilutions were plated in both KB with rifampicin and KB with rifampicin and carbenicillin agar plates. CFU of carbenicillin resistant colonies represent the Δ*exlx-gh5* strain. CFUs of Wt was determined as the count of total CFUs – carbenicillin resistant CFUs.

### Statistics

Statistical analyses were performed using Prism version 7.0 (GraphPad Software, La Jolla California USA, www.graphpad.com). Curves following initial symptoms in first leaf, systemic wilt in second leaf, and plant death from each experiment were compared using the built-in Log-rank (Mantel-Cox) test for survival analysis. In the cases where significant differences were found (*p* < 0.05), pairwise comparisons were tested using the same analysis (110). For comparisons of bacterial CFU *in planta*, CFU data and its log_10_-transformed values were checked for Gaussian distribution using the Shapiro-Wilk normality test. Since neither CFU data distribution nor the transformed Log_10_ values distribution passed the normality test, the Kruskal-Wallis non-parametric was used test to analyze if the medians vary significantly among experimental groups (*p* < 0.05). In the cases where differences were found, Dunn’s multiple comparisons test was used to test for pairwise differences between groups.

### Testing for cellulase and xylanase activity

Cellulase activity from cell-free supernatants of Wt and Δ*exlx-gh5* cultures were tested for extracellular enzymatic activity against cellulose. The Wt and Δ*exlx-gh5* strains were grown in 10 ml of liquid KB media for 48 H. Cultures were centrifuged at 7,000 rpm for 10 min, and each supernatant was filter-sterilized. Supernatants and 1 mg/ml cellulase (Sigma), were spotted in 1% agar, 1% Carboxy Methil Cellulose (CMC) plates. Plates were then incubated at 30°C for 48 h, and flooded with Gram’s Iodine. Halos were imaged after 24 h at RT.

A colony of *E. tracheiphila* grown on KB agar plates was used to test for extracellular xylanase activity. A xylanase producing strain of *Streptomyces lividens* was used as a positive control. Bacterial culture from each species was spotted on the surface of a KB agar plate, and grown at RT for 4 days. An overlay of 1% agar and 1% xylan was spread on top of the grown colonies, and plates were incubated at 30°C for 48 h. Plates were flooded with 1% congo red and incubated for 10 min before discarding the congo red solution. Plates were then flooded with 1N NaOH, and incubated for 10 min. NaOH was discarded and plates were imaged after 24 h at room temperature.

## Acknowledgements

This work was supported by Fundación Mexico en Harvard, and Conacyt grant 237414 to JR, NSF postdoctoral fellowship DBI-1202736 to LRS and NIH Grant GM58213 to RK. We thank all members of the Kolter lab, and Einat Segev, William R. Chase and Olga Zhaxybayeva for valuable feedback and discussion. We thank the staff at the Harvard Arnold Arboretum for assistance in plant cultivation, use of growth facilities and use of the confocal microscope. Dominique Schneider kindly donated the plasmid pDS132.

## References

1. Haichar FEZ, Marol C, Berge O, Rangel-castro J, Prosser JI, Balesdent J, Heulin T, Achouak W. 2008. Plant host habitat and root exudates shape soil bacterial community structure. The ISME Journal 2:1221–30.

2. Shi S, Richardson AE, O’Callaghan M, DeAngelis KM, Jones EE, Stewart A, Firestone MK, Condron LM. 2011. Effects of selected root exudate components on soil bacterial communities. FEMS Microbiology Ecology 77:600–610.

3. Lindow SE, Brandl MT. 2003. Microbiology of the Phyllosphere. Applied and Environmental Microbiology 69:1875–1883.

4. Huang X, Chaparro JM, Reardon KF, Zhang R. 2014. Rhizosphere interactions: root exudates, microbes and microbial communities. Botany.

5. Miwa H, Okazaki S. 2017. How effectors promote beneficial interactions. Current Opinion in Plant Biology 38:148–154.

6. Wees SCV, Ent SVd, Pieterse CM. 2008. Plant immune responses triggered by beneficial microbes. Current Opinion in Plant Biology 11:443–448.

7. Cosgrove DJ. 2017. Microbial Expansins. Annual Review of Microbiology 71:479–497.

8. Chase WR, Zhaxybayeva O, Rocha J, Cosgrove DJ, Shapiro LR. 2019. From morphogenesis to pathogenesis: A cellulose loosening protein is one of the most widely shared tools in nature. bioRxiv.

9. Georgelis N, Nikolaidis N, Cosgrove DJ. 2014. Biochemical analysis of expansin-like proteins from microbes. Carbohydrate Polymers 100:17–23.

10. Nikolaidis N, Doran N, Cosgrove DJ. 2013. Plant expansins in bacteria and fungi: evolution by horizontal gene transfer and independent domain fusion. Molecular Biology and Evolution 31:376–386.

11. Cosgrove DJ. 2000. Loosening of plant cell walls by expansins. Nature 407:321–326.

12. Sampedro J, Cosgrove DJ. 2005. The expansin superfamily. Genome biology 6:242.

13. Cosgrove DJ. 2015. Plant expansins: diversity and interactions with plant cell walls. Current Opinion in Plant Biology 25:162–172.

14. Chen F, Bradford KJ. 2000. Expression of an expansin is associated with endosperm weakening during tomato seed germination. Plant Physiology 124:1265–1274.

15. Brummell DA, Harpster MH, Civello PM, Palys JM, Bennett AB, Dunsmuir P. 1999. Modification of expansin protein abundance in tomato fruit alters softening and cell wall polymer metabolism during ripening. The Plant Cell 11:2203–2216.

16. Cho H-T, Cosgrove DJ. 2000. Altered expression of expansin modulates leaf growth and pedicel abscission in *Arabidopsis thaliana*. Proceedings of the National Academy of Sciences 97:9783–9788.

17. Li Y, Darley CP, Ongaro V, Fleming A, Schipper O, Baldauf SL, McQueen-Mason SJ. 2002. Plant expansins are a complex multigene family with an ancient evolutionary origin. Plant Physiology 128:854–864.

18. Cosgrove DJ, Li LC, Cho H-T, Hoffmann-Benning S, Moore RC, Blecker D. 2002. The growing world of expansins. Plant and Cell Physiology 43:1436–1444.

19. Georgelis N, Nikolaidis N, Cosgrove DJ. 2015. Bacterial expansins and related proteins from the world of microbes. Applied Microbiology and Biotechnology 99:3807–3823.

20. Kerff F, Amoroso A, Herman R, Sauvage E, Petrella S, Filée P, Charlier P, Joris B, Tabuchi A, Nikolaidis N, Cosgrove DJ. 2008. Crystal structure and activity of *Bacillus subtilis* YoaJ (EXLX1), a bacterial expansin that promotes root colonization. Proceedings of the National Academy of Sciences 105:16876–16881.

21. Kende H, Bradford K, Brummell D, Cho H-T, Cosgrove D, Fleming A, Gehring C, Lee Y, McQueen-Mason S, Rose J. 2004. Nomenclature for members of the expansin superfamily of genes and proteins. Plant Molecular Biology 55:311–314.

22. Saloheimo M, Paloheimo M, Hakola S, Pere J, Swanson B, Nyyssönen E, Bhatia A, Ward M, Penttilä M. 2002. Swollenin, a *Trichoderma reesei* protein with sequence similarity to the plant expansins, exhibits disruption activity on cellulosic materials. European Journal of Biochemistry 269:4202–4211.

23. Brotman Y, Briff E, Viterbo A, Chet I. 2008. Role of swollenin, an expansin-like protein from *Trichoderma*, in plant root colonization. Plant Physiology 147:779–789.

24. Tancos MA, Lowe-Power TM, Peritore-Galve FC, Tran TM, Allen C, Smart CD. 2018. Plant-like bacterial expansins play contrasting roles in two tomato vascular pathogens. Molecular Plant Pathology 19:1210–1221.

25. Chalupowicz L, Barash I, Reuven M, Dror O, Sharabani G, Gartemann KH, Eichenlaub R, Sessa G, Manulis-Sasson S. 2017. Differential contribution of *Clavibacter michiganensis* ssp. *michiganensis* virulence factors to systemic and local infection in tomato. Molecular Plant Pathology 18:336–346.

26. Laine MJ, Haapalainen M, Wahlroos T, Kankare K, Nissinen R, Kassuwi S, Metzler MC. 2000. The cellulase encoded by the native plasmid of *Clavibacter michiganensis* ssp. *sepedonicus* plays a role in virulence and contains an expansin-like domain. Physiological and Molecular Plant Pathology 57:221–233.

27. Hwang IS, Oh E-J, Lee HB, Oh C-S. 2019. Functional characterization of two cellulase genes in the Gram-positive pathogenic bacterium *Clavibacter michiganensis* for wilting in tomato. Molecular Plant-Microbe Interactions 32:491–501.

28. Hwang IS, Oh EJ, Kim D, Oh CS. 2018. Multiple plasmid-borne virulence genes of *Clavibacter michiganensis* ssp. *capsici* critical for disease development in pepper. New Phytologist 217:1177–1189.

29. Olarte-Lozano M, Mendoza-Nuñez MA, Pastor N, Segovia L, Folch-Mallol J, Martínez-Anaya C. 2014. PcExl1 a novel acid expansin-like protein from the plant pathogen *Pectobacterium carotovorum*, binds cell walls differently to BsEXLX1. PLoS One 9:e95638.

30. Jahr H, Dreier J, Meletzus D, Bahro R, Eichenlaub R. 2000. The endo-β-1, 4-glucanase CelA of Clavibacter michiganensis subsp. michiganensis is a pathogenicity determinant required for induction of bacterial wilt of tomato. Molecular Plant-Microbe Interactions 13:703–714.

31. Shapiro LR, Scully ED, Roberts D, Straub TJ, Geib SM, Park J, Stephenson AG, Rojas ES, Liu Q, Beattie G, Gleason M, Moraes CMD, Mescher MC, Fleischer SJ, Kolter R, Pierce N, Zhaxybayeva O. 2015. Draft genome sequence of Erwinia tracheiphila, an economically important bacterial pathogen of cucurbits. genomeA 3:e00482–15.

32. Shapiro LR, Andrade A, Scully ED, Rocha J, Paulson JN, Kolter R. 2018. Draft genome sequence of an Erwinia tracheiphila isolate from an infected muskmelon (Cucumis melo). Microbiology Resource Announcements 7:e01058–18.

33. . Shapiro L. 2012. A to ZYMV guide to Erwinia tracheiphila infection: An ecological and molecular study. The Pennsylvania State University.

34. Shapiro LR, Paulson JN, Arnold BJ, Scully ED, Zhaxybayeva O, Pierce N, Rocha J, Klepac-Ceraj V, Holton K, Kolter R. 2018. An introduced crop plant is driving diversification of the virulent bacterial pathogen *Erwinia tracheiphila*. mBio 9:e01307–18.

35. Shapiro LR, Mauck KE. 2018. Chemically-mediated Interactions among cucurbits, insects and microbes, p 55–90. In Tabata J (ed), Chemical Ecology of Insects. CRC Press.

36. . Smith EF. 1920. An introduction to bacterial diseases of plants. W.B. Saunders Company, Philadelphia.

37. Shapiro LR, Scully ED, Straub TJ, Park J, Stephenson AG, Beattie GA, Gleason ML, Kolter R, Coelho MC, Moraes CMD, Mescher MC, Zhaxybayeva O. 2016. Horizontal gene acquisitions, mobile element proliferation, and genome decay in the host-restricted plant pathogen *Erwinia tracheiphila*. Genome Biology and Evolution 8:649–664.

38. Moran NA, Plague GR. 2004. Genomic changes following host restriction in bacteria. Current Opinion in Genetics & Development 14:627–633.

39. de Mackiewicz D, Gildow FE, Blua M, Fleischer SJ, Lukezic FL. 1998. Herbaceous weeds are not ecologically important reservoirs of *Erwinia tracheiphila*. Plant Disease 82:521–529.

40. . Shapiro L, Seidl-Adams I, De Moraes C, Stephenson A, Mescher M. 2014. Dynamics of short-and long-term association between a bacterial plant pathogen and its arthropod vector. Scientific Reports 4.

41. Garcia-Salazar C, Gildow FE, Fleischer SJ, Cox-Foster D, Lukezic FL. 2000. ELISA versus immunolocalization to determine the association of *Erwinia tracheiphila* in *Acalymma vittatum* (Coleoptera: Chrysomelidae). Environmental Entomology 29:542–550.

42. Fleischer SJ, de Mackiewicz D, Gildow FE, Lukezic FL. 1999. Serological estimates of the seasonal dynamics of *Erwinia tracheiphila* in *Acalymma vittata* (Coleoptera: Chrysomelidae). Environmental Entomology 28:470–476.

43. Sasu M, Seidl-Adams I, Wall K, Winsor J, Stephenson A. 2010. Floral transmission of *Erwinia tracheiphila* by cucumber beetles in a wild *Cucurbita pepo*. Environmental Entomology 39:140–148.

44. Shapiro LR, Salvaudon L, Mauck KE, Pulido H, De Moraes CM, Stephenson AG, Mescher MC. 2013. Disease interactions in a shared host plant: effects of pre-existing viral infection on cucurbit plant defense responses and resistance to bacterial wilt disease. PLoS One 8:e77393.

45. Shapiro L, De Moraes CM, Stephenson AG, Mescher MC. 2012. Pathogen effects on vegetative and floral odours mediate vector attraction and host exposure in a complex pathosystem. Ecology Letters 15:1430–1438.

46. Sasu MA, Ferrari MJ, Du D, Winsor JA, Stephenson AG. 2009. Indirect costs of a nontarget pathogen mitigate the direct benefits of a virus-resistant transgene in wild *Cucurbita*. Proceedings of the National Academy of Sciences 106:19067–19071.

47. Marchler-Bauer A, Bo Y, Han L, He J, Lanczycki CJ, Lu S, Chitsaz F, Derbyshire MK, Geer RC, Gonzales NR. 2016. CDD/SPARCLE: functional classification of proteins via subfamily domain architectures. Nucleic Acids Research 45:D200–D203.

48. Lombard V, Golaconda Ramulu H, Drula E, Coutinho PM, Henrissat B. 2013. The carbohydrate-active enzymes database (CAZy) in 2013. Nucleic Acids Research 42:D490–D495.

49. Aziz R, Bartels D, Best A, DeJongh M, Disz T, Edwards R, Formsma K, Gerdes S, Glass E, Kubal M, Meyer F, Olsen G, Olson R, Osterman A, Overbeek R, McNeil L, Paarmann D, Paczian T, Parrello B, Pusch G, Reich C, Stevens R, Vassieva O, Vonstein V, Wilke A, Zagnitko O. 2008. The RAST Server: Rapid Annotations using Subsystems Technology. BMC Genomics 9:75.

50. Armenteros JJA, Tsirigos KD, Sønderby CK, Petersen TN, Winther O, Brunak S, von Heijne G, Nielsen H. 2019. SignalP 5.0 improves signal peptide predictions using deep neural networks. Nature Biotechnology 37:420–423.

51. Thomas CM, Nielsen KM. 2005. Mechanisms of, and barriers to, horizontal gene transfer between bacteria. Nature Reviews Microbiology 3:711.

52. Ochman H, Lawrence JG, Groisman EA. 2000. Lateral gene transfer and the nature of bacterial innovation. Nature 405:299–304.

53. Zhaxybayeva O, Doolittle WF. 2011. Lateral gene transfer. Current Biology 21:R242–R246.

54. Halpern M, Fridman S, Aizenberg-Gershtein Y, Izhaki I. 2013. Transfer of *Pseudomonas flectens* Johnson 1956 to *Phaseolibacter* gen. nov., in the family Enterobacteriaceae, as *Phaseolibacter flectens* gen. nov., comb. nov. International Journal of Systematic and Evolutionary Microbiology 63:268–273.

55. Mensi I, Vernerey M-S, Gargani D, Nicole M, Rott P. 2014. Breaking dogmas: the plant vascular pathogen *Xanthomonas albilineans* is able to invade non-vascular tissues despite its reduced genome. Royal Society Open Biology 4:130116.

56. Franklin NC. 1978. Genetic fusions for operon analysis. Annual Review of Genetics 12:193–221.

57. Vrisman CM, Deblais L, Rajashekara G, Miller SA. 2016. Differential colonization dynamics of cucurbit hosts by *Erwinia tracheiphila*. Phytopathology 106:684–692.

58. Cosgrove DJ. 2016. Catalysts of plant cell wall loosening. F1000Research 5.

59. Malinovsky FG, Fangel JU, Willats WG. 2014. The role of the cell wall in plant immunity. Frontiers in Plant Science 5:178.

60. Bahar O, Goffer T, Burdman S. 2009. Type IV pili are required for virulence, twitching motility, and biofilm formation of *Acidovorax avenae* subsp. *citrulli*. Molecular Plant-Microbe Interactions 22:909–920.

61. Burdman S, Bahar O, Parker JK, De La Fuente L. 2011. Involvement of type IV pili in pathogenicity of plant pathogenic bacteria. Genes 2:706–735.

62. Cursino L, Galvani CD, Athinuwat D, Zaini PA, Li Y, De La Fuente L, Hoch HC, Burr TJ, Mowery P. 2011. Identification of an operon, Pil-Chp, that controls twitching motility and virulence in Xylella fastidiosa. Molecular plant-microbe interactions 24:1198–1206.

63. Bahar O, Levi N, Burdman S. 2011. The cucurbit pathogenic bacterium *Acidovorax citrulli* requires a polar flagellum for full virulence before and after host-tissue penetration. Molecular Plant-Microbe Interactions 24:1040–1050.

64. Ewald PW. 1993. Evolution of infectious disease. Oxford University Press.

65. Mackinnon M, Read AF. 2004. Virulence in malaria: an evolutionary viewpoint. Philosophical Transactions of the Royal Society B: Biological Sciences 359:965–986.

66. Hinnebusch BJ, Joseph B, Perry RD, Schwan TG. 1996. Role of the *Yersinia pestis* hemin storage (hms) locus in the transmission of plague by fleas. Science 273:367.

67. Yao C, Zehnder, G., Bauske, E., Kloepper, J. 1996. Relationship between cucumber beetle (Coleoptera: Chrysomelidae) density and incidence of bacterial wilt of cucurbits. Journal of Economic Entomology 89:510–514.

68. Pasek S, Risler J-L, Brézellec P. 2006. Gene fusion/fission is a major contributor to evolution of multi-domain bacterial proteins. Bioinformatics 22:1418–1423.

69. Yang S BP. 2009. The evolutionary history of protein domains viewed by species phylogeny. PLoS ONE 4:e8378.

70. Boutilier MS, Lee J, Chambers V, Venkatesh V, Karnik R. 2014. Water filtration using plant xylem. PLoS One 9:e89934.

71. Pérez-Donoso AG, Sun Q, Roper MC, Greve LC, Kirkpatrick B, Labavitch JM. 2010. Cell wall-degrading enzymes enlarge the pore size of intervessel pit membranes in healthy and *Xylella fastidiosa*-infected grapevines. Plant Physiology 152:1748–1759.

72. Yadeta K, Thomma B. 2013. The xylem as battleground for plant hosts and vascular wilt pathogens. Frontiers in Plant Science 4.

73. Secchi F, Pagliarani C, Zwieniecki MA. 2017. The functional role of xylem parenchyma cells and aquaporins during recovery from severe water stress. Plant, Cell & Environment 40:858–871.

74. Christman MA, Sperry JS. 2010. Single-vessel flow measurements indicate scalariform perforation plates confer higher flow resistance than previously estimated. Plant, Cell & Environment 33:431–443.

75. Roper MC. 2011. *Pantoea stewartii* subsp. *stewartii*: lessons learned from a xylem-dwelling pathogen of sweet corn. Molecular Plant Pathology 12:628–637.

76. Foley JA, DeFries R, Asner GP, Barford C, Bonan G, Carpenter SR, Chapin FS, Coe MT, Daily GC, Gibbs HK. 2005. Global consequences of land use. Science 309:570–574.

77. Nasaruddin AS, Babler BN, Perna NT, Glasner JD, Charkowski AO. 2019. First report of *Dickeya dianthicola* causing blackleg on potato in Texas. Plant Disease.

78. Cai W, Srivastava S, Stulberg M, Nakhla M, Rascoe J. 2018. Draft genome sequences of two *Dickeya dianthicola* isolates from potato. Genome announcements 6:e00115–18.

79. Ma X, Schloop A, Swingle B, Perry KL. 2018. *Pectobacterium* and *Dickeya* responsible for potato blackleg disease in New York State in 2016. Plant Disease 102:1834–1840.

80. Dean R, Van Kan JA, Pretorius ZA, Hammond-Kosack KE, Di Pietro A, Spanu PD, Rudd JJ, Dickman M, Kahmann R, Ellis J. 2012. The ‘Top 10’ fungal pathogens in molecular plant pathology. Molecular Plant Pathology 13:414–430.

81. Mansfield J, Genin S, Magori S, Citovsky V, Sriariyanum M, Ronald P, Dow M, Verdier V, Beer SV, Machado MA. 2012. Top 10 plant pathogenic bacteria in molecular plant pathology. Molecular Plant Pathology 13:614–629.

82. Bae C, Han SW, Song Y-R, Kim B-Y, Lee H-J, Lee J-M, Yeam I, Heu S, Oh C-S. 2015. Infection processes of xylem-colonizing pathogenic bacteria: possible explanations for the scarcity of qualitative disease resistance genes against them in crops. Theoretical and Applied Genetics 128:1219–1229.

83. Agrios. 2005. Plant pathology, http://doc18.rupdfbook.com/plant-pathology-fifth-edition-by-george-n-agrios-PDFs-179089.pdf.

84. Metcalf R, Lampman R. 1989. The chemical ecology of Diabroticites and Cucurbitaceae. Experientia 45:240–247.

85. Joseph JG, Douglas WT, Edward GR, Anthony IC. 2008. Molecular phylogeny of rootworms and related galerucine beetles (Coleoptera: Chrysomelidae). Zoologica Scripta 37:195–222.

86. Rand FV, Cash LC. 1920. Some insect relations of *Bacillus tracheiphilus* Erw. Sm. Phytopathology 10:133–140.

87. Garcia-Salazar C, Gildow FE, Fleischer SJ, Cox-Foster D, Lukezic FL. 2000. Alimentary canal of adult *Acalymma vittata* (Coleoptera: Chrysomelidae): morphology and potential role in survival of *Erwinia tracheiphila* (Enterobacteriaceae). Canadian Journal of Entomology 132:1–13.

88. Roberts DC, Fleischer SJ, Sakamoto JM, Rasgon JL. 2018. Potential biological control of *Erwinia tracheiphila* by internal alimentary canal interactions in *Acalymma vittatum* with *Pseudomonas fluorescens*. Journal of Applied Microbiology 125:1137–1146.

89. Rand FV, Enlows EMA. 1916. Transmission and control of bacterial wilt of cucurbits. Journal of Agricultural Research 6:7–434.

90. Rand FV. 1915. Dissemination of bacterial wilt of cucurbits. Journal of Agricultural Research 5:257–260.

91. Whitaker TW, Bemis W. 1975. Origin and evolution of the cultivated Cucurbita. Bulletin of the Torrey Botanical Club:362–368.

92. Nee M. 1990. The domestication of *Cucurbita* (Cucurbitaceae). Economic Botany 44:56–68.

93. Schaefer H, Heibl C, Renner SS. 2009. Gourds afloat: a dated phylogeny reveals an Asian origin of the gourd family (Cucurbitaceae) and numerous oversea dispersal events. Proceedings of the Royal Society of London B: Biological Sciences 276:843–851.

94. Sebastian P, Schaefer H, Telford IRH, Renner SS. 2010. Cucumber (*Cucumis sativus*) and melon (*C. melo*) have numerous wild relatives in Asia and Australia, and the sister species of melon is from Australia. Proceedings of the National Academy of Sciences 107:14269–14273.

95. L.A. N, Trieu DA. 2011. Fusion gardens: Native North America and the Columbian Exchange. *In* Smith B (ed), Subsistence Economies of Indigenous North American Societies: A Handbook. Rowman and Littlefield Publishing Group, Lanham, MD.

96. Bisognin DA. 2002. Origin and evolution of cultivated cucurbits. Ciência Rural 32:715–723.

97. Datsenko KA, Wanner BL. 2000. One-step inactivation of chromosomal genes in *Escherichia coli* K-12 using PCR products. Proceedings of the National Academy of Sciences 97:6640–6645.

98. Altschul SF, Gish W, Miller W, Myers EW, Lipman DJ. 1990. Basic local alignment search tool. Journal of Molecular Biology 215:403–410.

99. Katoh K, Misawa K, Kuma Ki, Miyata T. 2002. MAFFT: a novel method for rapid multiple sequence alignment based on fast Fourier transform. Nucleic Acids Research 30:3059–3066.

100. Capella-Gutiérrez S, Silla-Martínez JM, Gabaldón T. 2009. trimAl: a tool for automated alignment trimming in large-scale phylogenetic analyses. Bioinformatics 25:1972–1973.

101. Abascal F, Zardoya R, Posada D. 2005. ProtTest: selection of best-fit models of protein evolution. Bioinformatics 21:2104–2105.

102. Stamatakis A. 2006. RAxML-VI-HPC: maximum likelihood-based phylogenetic analyses with thousands of taxa and mixed models. Bioinformatics 22:2688–2690.

103. Miller MA, Pfeiffer W, Schwartz T. Creating the CIPRES Science Gateway for inference of large phylogenetic trees, p 1–8. *In* (ed),

104. Sukumaran J, Holder MT. 2010. DendroPy: A Python library for phylogenetic computing. Bioinformatics 26:1569–1571.

105. Team RC. 2015. R: A language and environment for statistical computing. R Foundation of Statistical Computing Vienna, Austria.

106. Yu G, Smith DK, Zhu H, Guan Y, Lam TTY. 2017. ggtree: an R package for visualization and annotation of phylogenetic trees with their covariates and other associated data. Methods in Ecology and Evolution 8:28–36.

107. Guy L, Kultima JR, Andersson SG. 2010. genoPlotR: comparative gene and genome visualization in R. Bioinformatics 26:2334–2335.

108. Philippe N, Alcaraz J-P, Coursange E, Geiselmann J, Schneider D. 2004. Improvement of pCVD442, a suicide plasmid for gene allele exchange in bacteria. Plasmid 51:246–255.

109. Lagendijk EL, Validov S, Lamers GE, De Weert S, Bloemberg GV. 2010. Genetic tools for tagging Gram-negative bacteria with mCherry for visualization in vitro and in natural habitats, biofilm and pathogenicity studies. FEMS Microbiology Letters 305:81–90.

110. Machin D, Cheung YB, Parmar M. 2006. Survival analysis: a practical approach. John Wiley & Sons.

