## Supplemental for "A horizontally acquired expansin gene increases virulence of the emerging plant pathogen *Erwinia tracheiphila*"

**Running title*:*** *An expansin increases* Erwinia tracheiphila *virulence*

**Data Deposition Statement**: Analysis scripts and input files associated with reconstruction of phylogenetic trees are available at https://github.com/lshapiro31/gh5.expansin.phylogenetics

### **Supplemental Tables**

Supplemental Table 1. Strains and plasmids used in this study.

| Strains | Description | Reference |
|---|---|---|
| *Erwinia tracheiphila* BHKY | *Erwinia tracheiphila* parental Wild type strain. Spontaneous rifampicin resistant. Parental strain for mutants and complemented variants. | 32 |
| *Erwinia tracheiphila* BuffGH | Erwinia tracheiphila Wildtype isolate, used for fluorescence visualization of bacterial cells during infection. | |
| Δ*exlx-gh5* | Deletion mutant strain; *exlx-gh5:bla* genetic exchange. Ampicillin resistant. | This study |
| Δ*exlx* | Deletion mutant strain; clean deletion in N terminal coding region of *exlx*. | This study |
| Δ*eng* | Deletion mutant strain; *gh5:bla* genetic exchange. Ampicillin resistant. | This study |
| Δ*fliC* | Deletion mutant strain; *fliC*:*bla* genetic exchange. Ampicillin resistant. | This study |
| ΔT4P | Deletion mutant strain; *T4P*:*bla* genetic exchange. Ampicillin resistant. | This study |
| Δ*exlx-gh5*(cEXLX-GH5) | Mutant strain Δ*exlx-gh5,* complemented with wildtype promoter and *exp-gh5* coding region, inserted in a neutral chromosomic region. | This study |
| Δ*exlx*(EXLX) | Mutant strain Δ*exlx,* complemented with wildtype promoter and *exlx* coding region, inserted in a neutral chromosomic region. | This study |
| Δ*eng*(EXLX-ENG) | Mutant strain Δ*egh5,* complemented with wildtype promoter and *exlx-gh5* coding region, inserted in a neutral chromosomic region. | This study |
| *E. coli* Top10 | *E. coli* strain used for cloning. | |
| *E. coli* PIR1 | *E. coli* srtain for expression of R6K replication origin. | |
| *E. coli* S17-1λ | *E coli* strain used as donor for conjugation. | |
| Plasmids | | |
| pDS132 | Suicide plasmid for allelic replacement. | 108 |
| pMP7605 | Template for the amplification of the *mcherry* gene | 109 |
| pJR74 | Derived from pDS132, *mcherry* gene inserted in the *Xba*I site. | This study |
| pKD46 | Template for *bla* gene (ampicillin resistance cassete). | 97 |

| | | |
|---|---|---|
| pJR315 | Integration plasmid for *E. tracheiphila*. | This study |
| pJR150 | Plasmid carrying the construction for gene exchange between the *exlx-gh5* operon and the bla gene, derived from pJR74. | This study |
| pJR323 | Plasmid carrying the construction for *exlx* deletion, derived from pJR74. | This study |
| pJR324 | Plasmid carrying the construction for *gh5::bla* gene exchange, derived from pJR74. | This study |
| pJR74a | Plasmid carrying the construction for *fliC:bla* gene exchange, derived from pJR74. | This study |
| pJR149 | Plasmid carrying the construction for Type 4 Pili operon gene exchange with *bla* gene, derived from pJR74 | This study |
| pJR358 | Plasmid carrying the construction for *exlx-eng* operon integration into the chromosome of Et-BHKY, derived from pJR315. | This study |
| pJR357 | Plasmid carrying the construction for *exlx* gene integration into the chromosome of Et-BHKY, derived from pJR315. | This study |

Supplemental Table 2. Oligonucleotides used in this study.

| Primer Name | Description and sequence (5' to 3') | Modification |
|---|---|---|
| | ***bla* cassette amplification** | |
| LS23 | acttttcggggaaatgtgc | |
| LS24 | acgttaagggattttggtca | |
| | ***mcherry* gene amplification** | |
| JR72 | tcttctagacgttttcttacttgtacagctc | *Xba*I site |
| JR73 | tcttctagaaattcttgacaattaatcatcg | *Xba*I site |
| | ***exlx-gh5*::*bla* exchange** | |
| **LS52 (Fwd 5')** | acccgttaatgcaccagaac | |
| **LS53 (reamplification)** | gaggagctcttatttcgatgatggttttatgg | *Sac*I site |
| **LS54 (Rev 5')** | gcacatttccccgaaaagtagttaaaacagcgcagatgg | LS23 tag |
| **LS55 (Fwd 3')** | tgaccaaaatcccttaacgtacagaggatgccctggtaag | LS24 tag |
| **LS57 (reamplification)** | gaggagctccgcaaatcatcaccagtcag | *Sac*I site |
| **LS56 (Rev 3')** | gaggtatatcccggcctgac | |
| | ***exlx* clean deletion** | |
| **JR175 (Fwd 5')** | Cagaactgacgcttacctcc | |

| | | |
|---|---|---|
| **JR176 (reamplification)** | GAGGAGCTCtcgcttctttatagagctgc | *Sac*I site |
| JR194 (Rev 5') | gttgagctggtaaggactaagg gaaagtcagagccgctattG | JR195 tag |
| JR195 (Fwd 3') | Caatagcggctctgactttcccttagtccttaccagctcaac | JR194 tag |
| **JR179 (reamplification)** | GAGGAGCTCatcgataacgctatccacac | *Sac*I site |
| **JR180 (Rev 3')** | Gtatttcgtagccaatctctg | |
| | ***gh5::bla* exchange** | |
| **JR181 (Fwd 5')** | Gaatcaggcagacttggttc | |
| **JR182 (reamplification)** | GAGGAGCTCgtgattacaaccatgctttcg | *Sac*I site |
| **JR183 (Rev 5')** | GCACATTTCCCCGAAAAGT gcatgctgctacagataacc | LS23 tag |
| **JR184 (Fwd 3')** | TGACCAAAATCCCTTAACGT gataaagcagaaggagcatc | LS24 tag |
| **JR185 (reamplification)** | GAGGAGCTCcccactgatgttatcgctc | *Sac*I site |
| **JR186 (Rev 3')** | Catgctgtttttttatattacctgc | |
| | **Integration plasmid** | |
| **JR143 (Fwd 5')** | GAGGAGCTCcttcaaaaatacgttcacacc | *Sac*I site |
| **JR144 (Rev 5')** | gttcatacatggtcaaacctg AGATCTCTCGAG catttatccgtgctgatctg | *Xho*I, *Bgl*II sites, JR145 tag |
| **JR145 (Fwd 3')** | cagatcagcacggataaatg CTCGAGAGATCT caggtttgaccatgtatgaac | *Xho*I, *Bgl*II sites, JR144 tag |
| **JR146 (Rev 3')** | GAGGAGCTCaataccaatcaggaccacac | *Sac*I site |
| | **Complementation of *exlx-gh5* and *exlx*** | |
| **JR152 (Fwd)** | CTCCTCGAGggaagattttcatcagcacc | *Xho*I site |
| **JR153 (Rev *exlx* only)** | CTCCTCGAGctgtcatgtcctgtattatatattgtg | *Xho*I site |
| **JR154 (Rev *exlx-eng*)** | CTCCTCGAGgacagtaccagtatcctgacg | *Xho*I site |

Supplemental Table 3. Species and sequences included in Figure 2.

| Species | Location | Accession | Description |
|---|---|---|---|
| ***Erwinia tracheiphila*** | Chromosome | WP_046372116.1, WP_016193008.1 | Expansin - GH5 locus |
| ***Pantoea stewartii*** | Plasmid pDSJ08 | WP_044243227.1, WP_006122111.1 | Expansin - GH5 locus |
| ***Pectobacterium carotovorum*** | Chromosome | WP_103860671.1 | Single expansin domain |
| ***Dickeya solani*** | Chromosome | WP_022633560.1 | Single expansin domain |

| | | | |
|---|---|---|---|
| ***Dickeya zeae*** | Chromosome | WP_026358096.1 | Single expansin domain |
| ***Dickeya dianthicola*** | Chromosome | WP_024104808.1, WP_103415919.1 | Expansin - GH5 locus |
| ***Lonsdalea quercina*** | Chromosome | WP_026740116.1 | Single expansin domain |
| ***Lonsdalea iberica*** | | WP_094110348.1 | Single expansin domain |
| ***Lonsdalea britannica*** | Chromosome | WP_094118757.1 | Single expansin domain |
| ***Brennaria salicis*** | | WP_113865163.1 | Single expansin domain |
| ***Pectobacterium wasabiae*** | Chromosome | WP_005968506.1 | Single expansin domain |
| ***Pectobacterium betavasculorum*** | Chromosome | WP_010275007.1 | Single expansin domain |
| ***Pectobacterium peruviense*** | Chromosome | WP_113627469.1 | Single expansin domain |
| ***Brenneria salicis*** | Chromosome | WP_113865163.1 | Single expansin domain |
| ***Xanthomonas sacchari*** | Chromosome | WP_043094747.1 | Single expansin domain |
| ***Xanthomonas albilineans*** | Chromosome | WP_045769814.1 | Single expansin domain |
| ***Xanthomonas translucens*** | Chromosome | WP_003477706.1 | Single expansin domain |
| ***Xanthomonas arboricola*** | Chromosome | SUZ34847.1 | Single expansin domain |
| ***Xanthomonas maliensis*** | Chromosome | WP_031340658.1 | Fusioned 1,4-beta-cellobiosidase-expansin |
| ***Xylella fastidiosa*** | Chromosome | WP_020851755.1 | Fusioned 1,4-beta-cellobiosidase-expansin |
| ***Xylella taiwanensis*** | Chromosome | WP_038269744.1 | Fusioned 1,4-beta-cellobiosidase-expansin |
| ***Xanthomonas oryzae*** | Chromosome | WP_131078740.1 | Fusioned 1,4-beta-cellobiosidase-expansin |
| ***Xanthomonas citri*** | Chromosome | WP_096035367.1 | Fusioned 1,4-beta-cellobiosidase-expansin |
| ***Xanthomonas campestri*** | Chromosome | KFA09681.1 | Fusioned 1,4-beta-cellobiosidase- |

| | | | |
|---|---|---|---|
| | | | expansin |
| ***Xanthomonas phaseoli*** | Chromosome | WP_017161821.1 | Fusioned 1,4-beta-cellobiosidase-expansin |
| ***Ralstonia pseudosolanacearum*** | Chromosome | WP_120452416.1 | Single expansin domain |
| ***Ralstonia solanacearum*** | Chromosome | WP_134927564.1 | Single expansin domain |
| ***Myxococcus virescens*** | Chromosome | WP_090491481.1 | Single expansin domain |
| ***Myxococcus fulvus*** | Chromosome | WP_046714021.1 | Single expansin domain |
| ***Clavibacter michiganensis*** | Plasmid pCM1 | PRJNA19643, AM711865.1 | Multidomain GH5-CBMII-expansin, |
| | Chromosome | 4JCW_A | Single expansin domain |
| ***Streptomyces scabiei*** | Chromosome | FN554889.1, WP_013005169.1 | Single expansin domain |
| | Chromosome | CBG75940.1 | Single expansin domain |
| ***Streptomyces cellulosae*** | Chromosome | WP_030660953.1 | SSL4- expansin fusion |
| ***Streptomyces griseus*** | Chromosome | WP_051866281.1 | Single expansin domain |
| ***Streptomyces ipomoeae*** | Chromosome | WP_009327266.1 | Single expansin domain |
| ***Paenibacillus xylanexedens*** | Chromosome | WP_124115839.1 | Single expansin domain |
| ***Bacillus licheniformis*** | Chromosome | WP_073425687.1 | Single expansin domain |
| ***Bacillus glycinifermentans*** | Chromosome | WP_048355312.1 | Single expansin domain |
| ***Bacillus pumilus*** | Chromosome | WP_024718945.1 | Single expansin domain |
| ***Bacillus cellulasensis*** | Chromosome | APP16610.1 | Single expansin domain |
| ***Bacillus halotolerans*** | Chromosome | WP_106021706.1 | Single expansin domain |
| ***Bacillus subtilis*** | Chromosome | WP_128737925.1 | Single expansin domain |
| ***Acidovorax avenae*** | Chromosome | WP_107177641.1 | Fusioned CelA1 cellulase - expansin |
| ***Acidovorax radicis*** | Chromosome | WP_010459029.1 | Single expansin |

| | | | |
|---|---|---|---|
| | | | domain |
| ***Acidovorax citruli*** | Chromosome | ABM33954.1 | Fusioned CelA1 cellulase - expansin |

**Supplemental Figures and Legends**

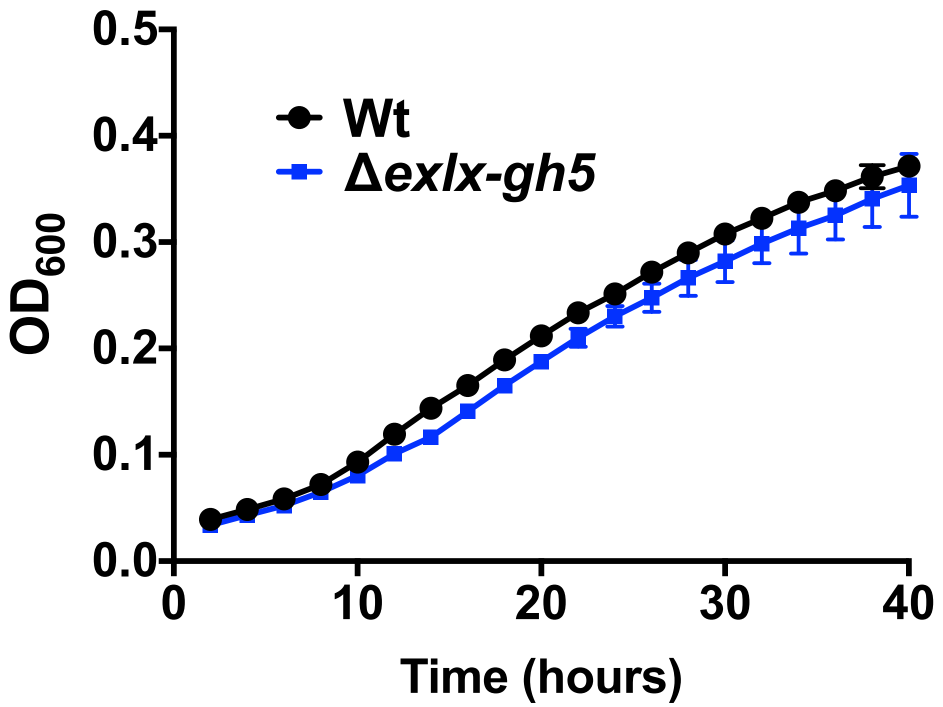


**Supplemental Figure 1. *In vitro* growth of Wt and Δ*exlx-gh5*.** A single colony of Wt or Δ*exlx-gh5* strains were picked into 3 ml of liquid Kb media and grown for 48 h at 25°C with shaking. Then, 1 ml of each culture was washed once with 1 volume of PBS and diluted with fresh KB media to $OD_{600}$ of 0.05. Four replicates of 300 µl were placed in a clear 96-well microplate and growth was followed in these standing liquid cultures using a microplate reader, by measuring absorbance at 600 nm every 2 h for 40 h at a constant temperature of 25°C. Average ± SD is shown.

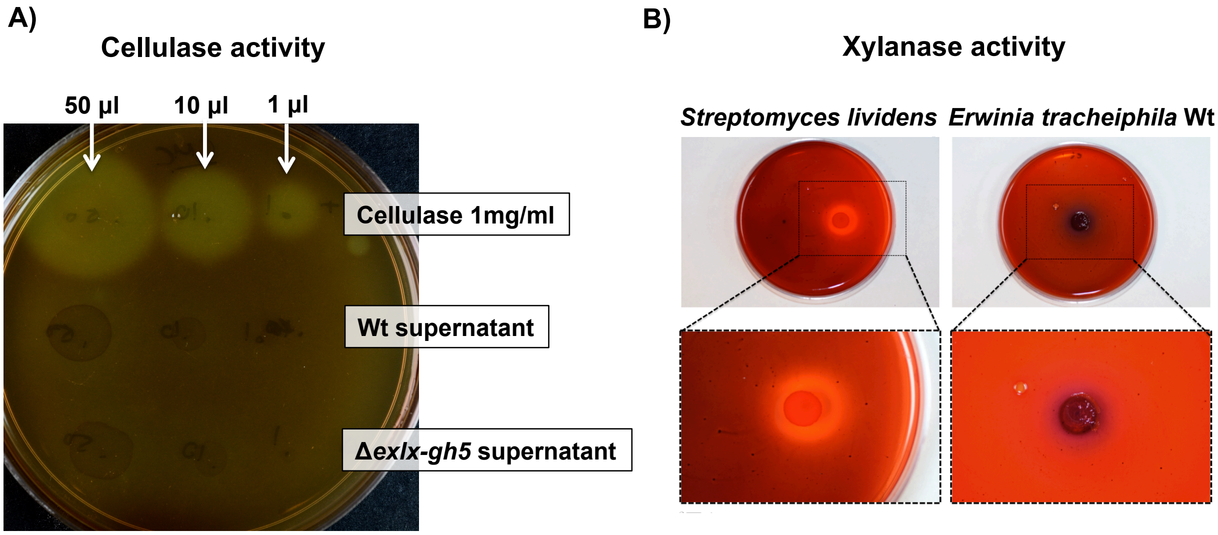


**Supplemental Figure 2**. **Tests to detect *E. tracheiphila* cellulase and xylanase activity *in vitro*. A)** Commercial cellulase and supernatants of Wt and the Δ*exlx-gh5* mutant were spotted in media containing agar and 1% CMC, incubated at 30°C for 48 h, and then flooded with Gram's Iodine. Halos were imaged after 24 h at RT. **B)** *E. tracheiphila* and the xylan degrading species *Streptomyces lividens* were grown in KB agar. An overlay of 1% xylan and 1% agar was spread on top of the colonies. Plates were incubated at 30°C for 48 h, and flooded with 1% Congo Red. Halos were imaged and measured after 24 h at room temperature.
